# Regulation of Neural Circuit Development by Cadherin-11 Provides Implications for Autism

**DOI:** 10.1101/2020.04.24.058438

**Authors:** Jeannine A. Frei, Robert F. Niescier, Morgan S. Bridi, Madel Durens, Jonathan E. Nestor, Michaela B. C. Kilander, Xiaobing Yuan, Derek M. Dykxhoorn, Michael W. Nestor, Shiyong Huang, Gene J. Blatt, Yu-Chih Lin

**Author notes:** Correspondence: Jeannine A. Frei Yu-Chih Lin.

## Abstract

Autism spectrum disorder (ASD) is a neurological condition characterized by alterations in social interaction and communication, and restricted and/or repetitive behaviors. The classical type II cadherins cadherin-8 (Cdh8, CDH8) and cadherin-11 (Cdh11, CDH11) have been implicated as autism risk gene candidates. To explore the role of cadherins in the etiology of autism, we investigated their expression patterns during mouse brain development and in autism-specific human tissue. In mice, expression of cadherin-8 and cadherin-11 was developmentally regulated and enriched in the cortex, hippocampus, and thalamus/striatum during the peak of dendrite formation and synaptogenesis. Both cadherins were expressed in synaptic compartments but only cadherin-8 associated with the excitatory synaptic marker neuroligin-1. Induced pluripotent stem cell (iPSC)-derived cortical neural precursor cells (NPCs) and cortical organoids generated from individuals with autism showed upregulated CDH8 expression levels while CDH11 expression levels were downregulated. We used *Cdh11* knockout mice of both sexes to analyze the function of cadherin-11, which could help explain phenotypes observed in autism. *Cdh11^-/-^* hippocampal neurons exhibited increased dendritic complexity along with altered neuronal and synaptic activity. Similar to the expression profiles in human tissue, levels of cadherin-8 were significantly elevated in *Cdh11* knockout brains. Additionally, excitatory synaptic markers neuroligin-1 and PSD-95 were both increased. Together, these results strongly suggest that cadherin-11 is involved in regulating the development of neuronal circuitry and that alterations in the expression levels of cadherin-11 may contribute to the etiology of autism.

**Significance Statement:** Autism is a neurodevelopmental condition with high genetic and phenotypic heterogeneity. Multiple genes have been implicated in autism, including the cadherin superfamily of adhesion molecules, cadherin-8 and cadherin-11. This study first characterizes the expression profiles of cadherin-8 and cadherin-11 to understand the potential roles they play in the development of neurons. The study further describes novel contributions of cadherin-11 in neural circuit formation. Loss of cadherin-11 in mice results in altered levels of several synaptic proteins, including PSD-95, neuroligin-1, and cadherin-8, and changes the morphology and activity of excitatory neurons. The levels of cadherin-8 and cadherin-11 in human cells of autistic individuals are both altered, strengthening the hypothesis that these two cadherins may involve in aspects of autism etiology.

## Introduction

Autism is a neurodevelopmental condition characterized by marked qualitative changes in social interaction and communication as well as restricted and repetitive behaviors (American Psychiatric Association, 2013). Current estimates indicate that 1 in 54 children in the United States are affected (Maenner et al., 2020). Autism is characterized by both phenotypic and genetic heterogeneity. This genetic complexity is illustrated by the fact that no single gene associated with the condition contributes to more than 1% of the autism cases (Huguet et al., 2013; Jeste and Geschwind, 2014). Nevertheless, many of the genes implicated in the condition encode for synaptic cell adhesion molecules, scaffolding proteins and cytoskeletal regulators (Betancur et al., 2009; Bourgeron, 2015; Lin et al., 2016). These autism risk genes converge into selective cellular pathways that appear to be commonly affected, including neurite outgrowth, dendritic spine stability, synaptogenesis, and synaptic function (Betancur et al., 2009; Hussman et al., 2011; Chen et al., 2014; Bourgeron, 2015; Lin et al., 2016; Joensuu et al., 2017).

One large group of synaptic cell adhesion molecules, the cadherin superfamily, has been widely associated with neurodevelopmental disorders, including autism (Redies et al., 2012; Lin et al., 2016). The cadherin family comprises more than one hundred members, which are further divided into subfamilies, including classical type I and II cadherins, clustered and non-clustered protocadherins, and atypical FAT cadherins (Hulpiau and van Roy, 2009; Hirano and Takeichi, 2012). Multiple studies have identified cadherins across all subfamilies as candidate risk genes for autism (Marshall et al., 2008; Morrow et al., 2008; Depienne et al., 2009; Wang et al., 2009; Willemsen et al., 2010; Chapman et al., 2011; Hussman et al., 2011; Pagnamenta et al., 2011; Sanders et al., 2011; Camacho et al., 2012; Neale et al., 2012; O’Roak et al., 2012; Girirajan et al., 2013; van Harssel et al., 2013; Crepel et al., 2014; Cukier et al., 2014; Kenny et al., 2014).

The most-well studied classical type I cadherin, N-cadherin, is broadly expressed in the central nervous system and has been implicated in multiple processes during nervous system development (Takeichi and Abe, 2005; Arikkath and Reichardt, 2008; Hirano and Takeichi, 2012; Friedman et al., 2015a; Seong et al., 2015). In contrast to classical type I cadherins, the expression profile of classical type II cadherins is more restricted to specific brain circuits and subcellular compartments (Korematsu and Redies, 1997; Suzuki et al., 1997; Inoue et al., 1998; Wang et al., 2019). The differential expression patterns of classical type II cadherins allow them to confer sophisticated synaptic specificity (Basu et al., 2015).

The aim of the present study was to investigate the involvement of classical type II cadherins in autism and to discern the contribution of these cadherins to neuronal development and function. We focused our study on the classical type II cadherins cadherin-8 (Cdh8, CDH8) and cadherin-11 (Cdh11, CDH11) as these specific cadherins have been identified as autism risk genes in a genome-wide association study and by whole exome sequencing (Hussman et al., 2011; Cukier et al., 2014). In addition, other studies have identified rare mutations and SNP variants in *CDH8* and *CDH11* genes, respectively, in individuals with autism (Pagnamenta et al., 2011; Crepel et al., 2014). Here, we first examined the expression profiles and binding partners of cadherin-8 and cadherin-11 in developing mouse brains. Our results indicated that although these two cadherins show similar spatial expression and interact with each other, they may engage in divergent cellular pathways. We used *Cdh11* knockout mice to identify potential phenotypes that may be altered in autism as, similarly, human induced pluripotent stem cell (hiPSC)-derived neural precursor cells (NPCs) and organoids generated from individuals with autism displayed differentially altered expression levels of cadherin-8 and cadherin-11. Together, the data suggest that depletion of cadherin-11 causes altered neural circuit development that may drive aspects of autism pathophysiology.

## Materials and Methods

### Animals

C57BL/6 mice were purchased from the animal facility of the University of Maryland School of Medicine Program in Comparative Medicine (Baltimore, MD, USA). *Cdh11^tm1Mta^/*HensJ mice were purchased from the Jackson Lab (Horikawa et al., 1999). Mice were housed and cared for by the AAALAC accredited program of the University of Maryland School of Medicine. Female mice were group-housed and male mice were singly housed with *ad libitum* food and water accessibility under a standard 12-hour light/dark cycle. Neonatal mice of both sexes were euthanized for the preparation of neuronal and non-neuronal cultures. To match the mixed-gender condition in cultures animals of both sexes were used for biochemistry. All experiments were reviewed and approved by the Institutional Care and Use Committees (IACUC) of the University of Maryland School of Medicine and the Hussman Institute for Autism, and were performed in accordance with the animal care guidelines of the National Institute of Health.

### Antibodies

Primary and secondary antibodies used in this study are listed in Table 1 and Table 2, respectively. The specificity of the antibodies was carefully examined prior to conducting the experiments (Figure 1-1).

**Figure 1.**
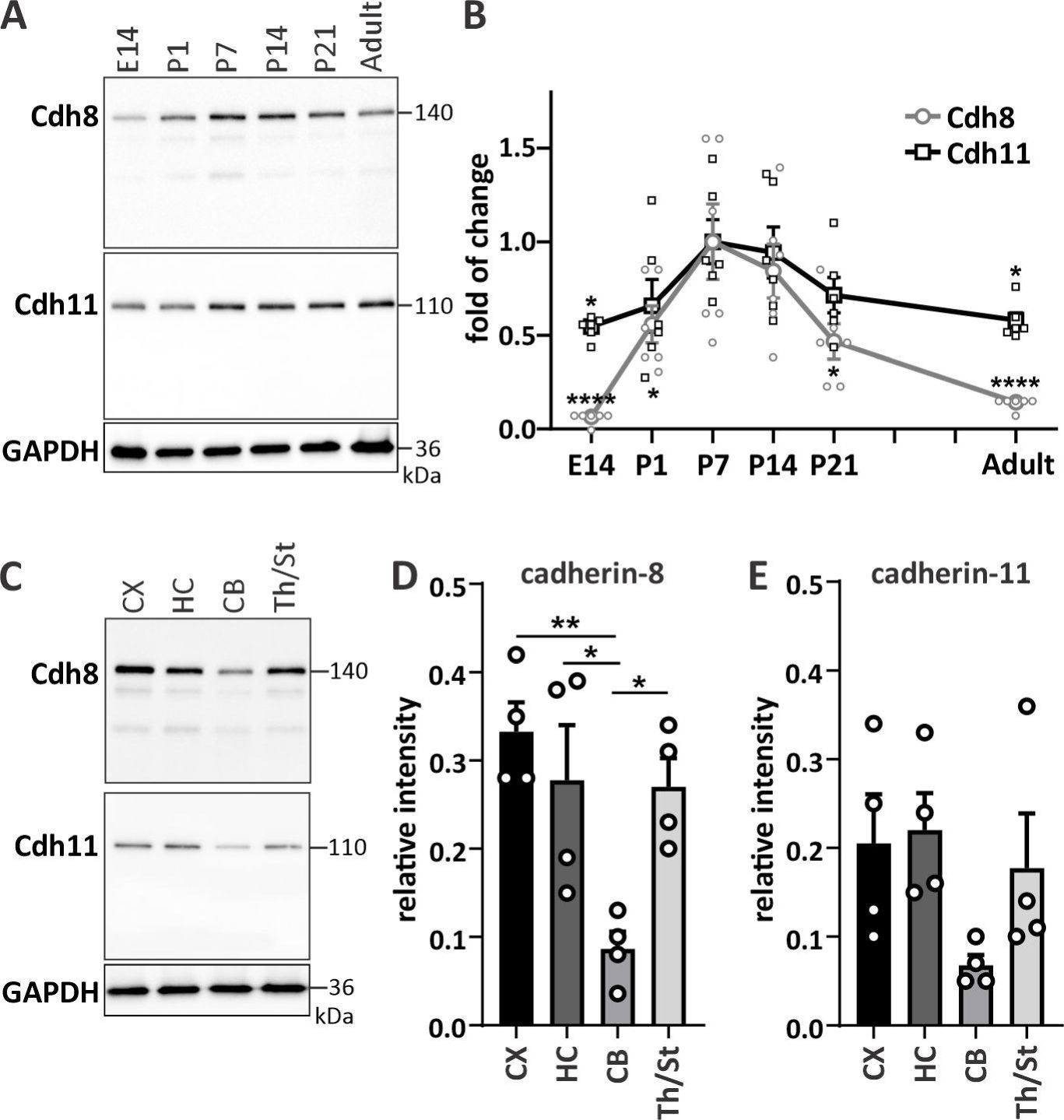
Cadherin-8 and cadherin-11 show similar temporal and spatial expression patterns. **(A)** Temporal expression profile of cadherin-8 and cadherin-11 in mouse whole brain harvested at different developmental ages. Adult mice were 5 months old. **(B)** Line graph of temporal expression of the two cadherins. Values were normalized to P7. Cdh8: \**p* = 0.046 (P1), \**p* = 0.0123 (P21), \*\*\*\**p* < 0.0001; Cdh11: \**p* = 0.017 (E14), \**p* = 0.0296 (adult), one-way ANOVA with Dunnett’s multiple comparisons test. N = 6 whole brains per age from three independent litters. **(C)** Expression profiles of cadherin-8 and cadherin-11 in different brain areas, including cortex (CX), hippocampus (HC), cerebellum (CB) and thalamus/striatum (Th/St) at P14. Quantification of **(D)** cadherin-8 and **(E)** cadherin-11 expression in different brain regions. \**p* = 0.0259 between HC and CB, \**p* = 0.0326 between CB and Th/St, \*\**p* = 0.0048 between CX and CB, one-way ANOVA with Tukey’s multiple comparisons test. N = 4 samples per brain area with 2-3 pooled brain areas per sample. Cadherin signals were normalized to GAPDH.

**Table 1.**
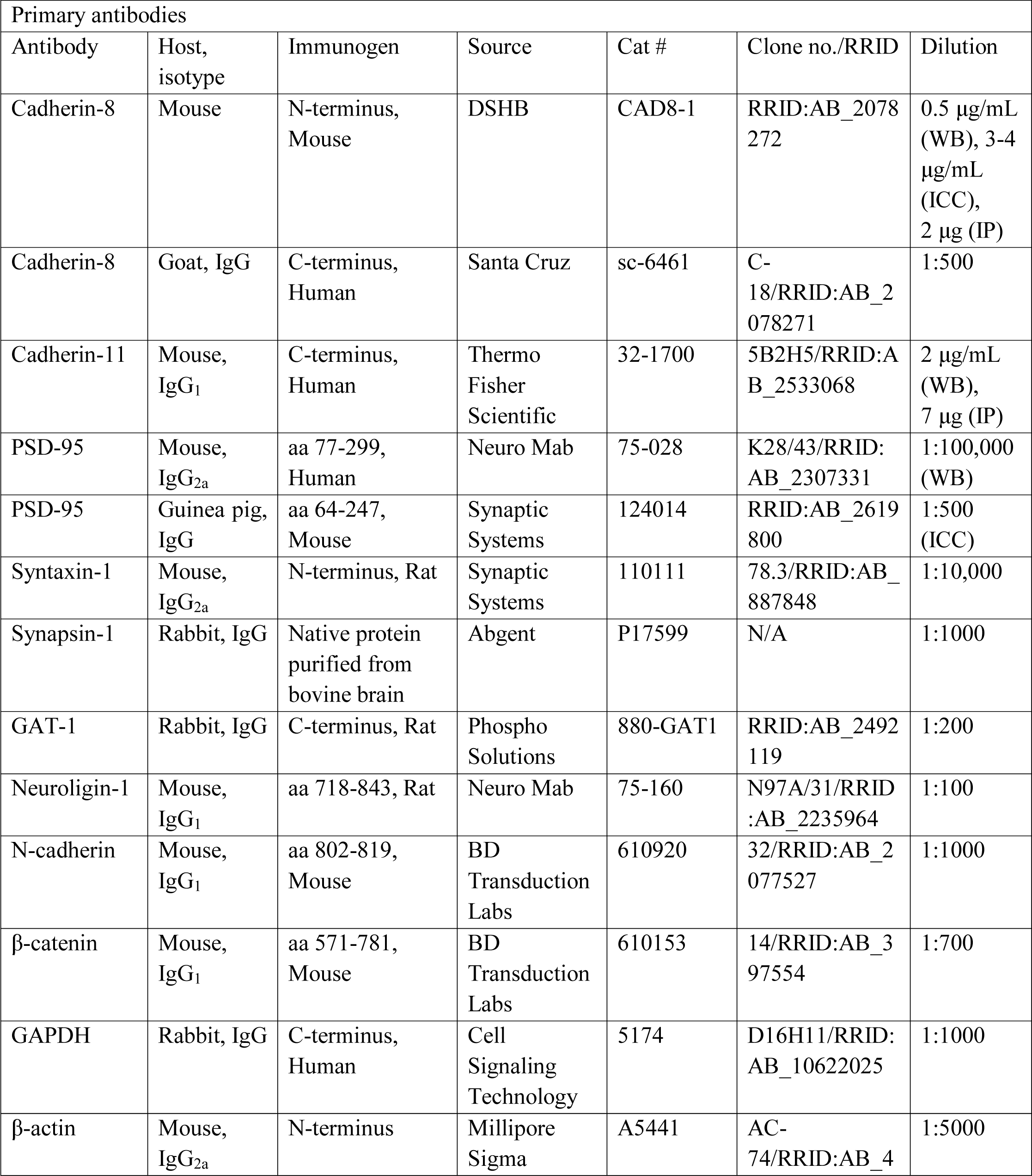
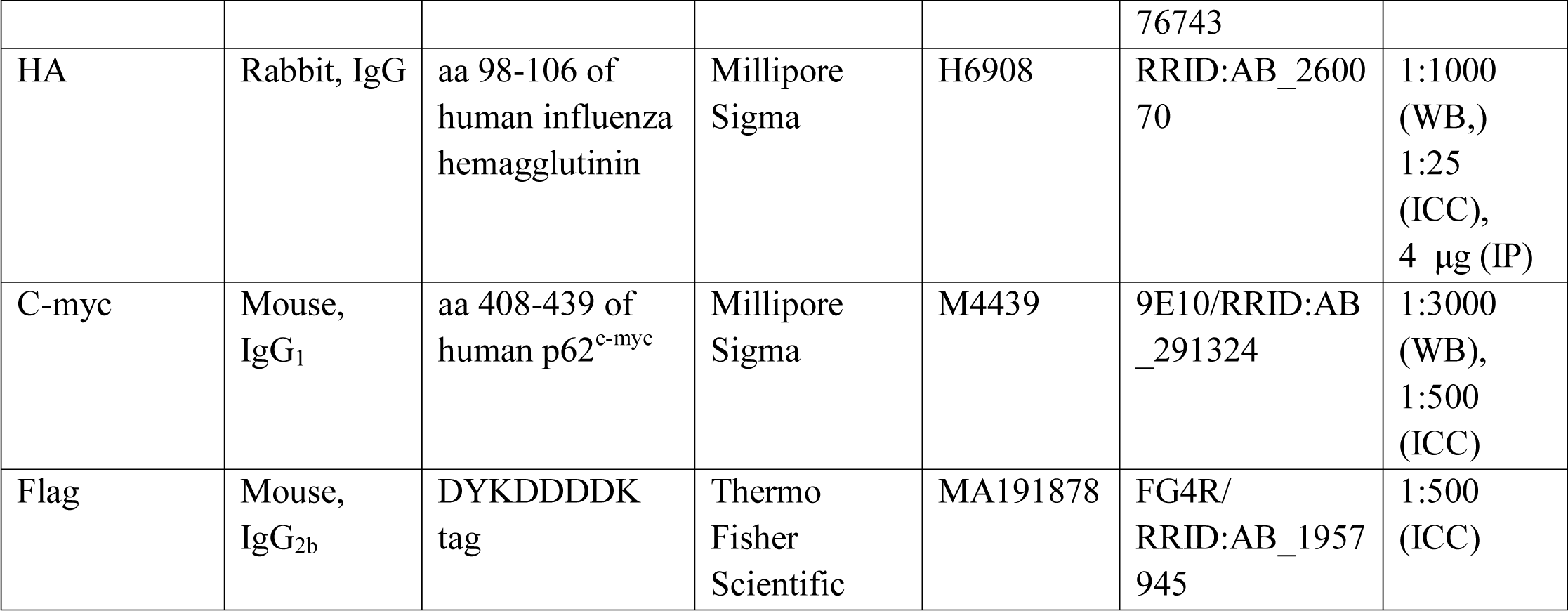
Primary antibodies. List of primary antibodies used for Western blot (WB), immunocytochemistry (ICC) and immunoprecipitation (IP). Abbreviations: PSD-95, postsynaptic density protein-95; GAT-1, sodium- and chloride-dependent GABA transporter-1; GAPDH, glyceraldehyde 3-phosphate dehydrogenase; aa, amino acid.

**Table 2.** Secondary antibodies. List of secondary antibodies used for Western blot and immunocytochemistry. Abbreviations: HRP, horseradish peroxidase.

| Secondary antibodies |  |  |  |  |  |  |
| --- | --- | --- | --- | --- | --- | --- |
| Antibody | Host, isotype | Immunogen | Source | Cat # | Clone no./RRID | Dilution |
| anti-rabbit IgG, HRP conjugate | Goat, IgG | IgG Rabbit | Cell Signaling Technology | 7074P2 | RRID:AB_2099233 | 1:7500 |
| anti-mouse IgG, HRP conjugate | Goat, IgG | IgG Mouse | Thermo Fisher Scientific | A16072 | RRID:AB_2534745 | 1:5000 |
| anti-goat IgG, HRP conjugate | Donkey, IgG | IgG Goat | Thermo Fisher Scientific | A15999 | RRID:AB_2534673 | 1:5000 |
| anti-mouse Alexa488 | Donkey, IgG | IgG (H+L) Mouse | Thermo Fisher Scientific | A21202 | RRID:AB_141607 | 1:1000 |
| Anti-mouse Alexa647 | Donkey, IgG | IgG (H+L) Mouse | Thermo Fisher Scientific | A31571 | RRID:AB_162542 | 1:1000 |
| anti-rabbit Alexa488 | Donkey, IgG | IgG (H+L) Rabbit | Thermo Fisher Scientific | A21206 | RRID:AB_2535792 | 1:1000 |
| anti-rabbit Alexa568 | Donkey, IgG | IgG (H+L) Rabbit | Thermo Fisher Scientific | A10042 | RRID:AB_2534017 | 1:1000 |
| anti-rabbit Alexa647 | Donkey, IgG | IgG (H+L) Rabbit | Thermo Fisher Scientific | A31573 | RRID:AB_2536183 | 1:1000 |
| anti-guinea pig Alexa488 | Donkey, IgG | IgG (H+L) Guinea Pig | Jackson Immuno Research | 706545148 | RRID:AB_2340472 | 1:1000 |
| anti-guinea pig Alexa568 | Goat, IgG | IgG (H+L) Guinea Pig | Thermo Fisher Scientific | A11075 | RRID:AB_2534119 | 1:1000 |

### Plasmids

Myc-flag-tagged full-length *Cdh8* was purchased from Origene (plasmid #MR218916). *Cdh8* was expressed under the CMV promoter in the pCMV6 vector. Flag-tagged full-length *Cdh11* was expressed under the EF-1α promoter in the pBos vector (gift from Dr. Megan Williams, University of Utah, USA). HA-tagged *Nlgn1* plasmid was a gift from Peter Scheiffele (Addgene plasmid # 15260; RRID:Addgene_15260) (Chih et al., 2006). *Nlgn1* was expressed under the chicken β-actin promoter in the pCAAGs vector. pLL3.7-GFP was a gift from Luk Parijs β (Addgene plasmid # 11795; RRID:Addgene_11795) (Rubinson et al., 2003).

### Cell cultures and transfection

Non-neuronal cell cultures were prepared from postnatal day 0 (P0) C57BL/6 mouse cortices and cultured in Dulbecco’s MEM growth medium (Invitrogen Cat#11960044) supplemented with 10% FBS (Millipore Sigma Cat#F4135), 2 mM L-glutamine (Invitrogen Cat#25030081) and 1% penicillin/streptomycin (Invitrogen Cat#15140122). Primary neuronal cultures were prepared from P0 C57BL/6 mouse cortex (3-4 animals per culture) or hippocampus (8-10 animals per culture). Hippocampal cultures from *Cdh11^tm1Mta^/*HensJ mice were prepared from individual pups at P0. The genotype of each pup was determined after the culture was prepared. Only *Cdh11* wild-type and knockout cultures were used for further experiments. In brief, brain tissue was dissected and meninges were removed. Tissue was digested in papain and cells were dissociated and plated on surfaces coated with 20 μg/ml poly-D-lysine (Millipore Sigma μ Cat#P6407). Cortical and hippocampal cultures were maintained in serum-free Neurobasal-A media (Invitrogen Cat#10888022) containing 2 mM L-glutamine (Gibco Cat#25030081), 1% penicillin/streptomycin (Gibco Cat#15140122) and 2% B27 supplement (Gibco Cat#17504044). For Western blot analysis, non-neuronal cells were harvested at 14 days *in vitro* (DIV) and cortical neurons were harvested at different time points: 1, 3, 7 and 14 DIV. *Cdh11* wild-type and knockout cultures were harvested at 4, 7 and 14 DIV. Neuro-2A (N2a, mouse neuroblastoma cell line; ATCC) cells were maintained in Dulbecco’s MEM growth medium (Invitrogen Cat#11960044) supplemented with 10% FBS (Millipore Sigma Cat#F4135), 2 mM L-glutamine (Gibco Cat#25030081) and 1% penicillin/streptomycin (Gibco Cat#15140122). N2a cells were transfected with full-length plasmids using Lipofectamine 3000 (Invitrogen Cat#L3000015) according to the manufacturer’s protocol. Cells were harvested 48 hours post-transfection.

### Western blot analysis

At the indicated points in development, mice of both sexes were sacrificed and brains were quickly dissected. Either whole brains or different brain areas, including forebrain, cortex, hippocampus, cerebellum and thalamus/striatum were collected. Brain tissue was snap-frozen in liquid nitrogen. All tissues and cells were lysed in RIPA buffer (Cell Signaling Technologies Cat# 9806S) supplemented with PMSF (Cell Signaling Technologies Cat#8553S) and protease and phosphatase inhibitor cocktail (Thermo Fisher Scientific Cat#78442). Protein concentration was determined using Pierce BCA protein assay kit (Thermo Fisher Scientific Cat#23227) and measured by the Tecan Spark 10M multimode microplate reader. Ten micrograms of protein samples from brain or cell lysates were run on 10% Tris-glycine SDS-PAGE gels and transferred to a PVDF membrane. Membranes were blocked in 5% milk/TBS-T followed by incubation of primary antibodies overnight and secondary HRP-coupled antibodies for one hour at room temperature. Blots were imaged with the ChemiDoc Touch Imaging System (Bio Rad) and band densitometries were analyzed using the Image Lab software (Bio Rad). Intensities were normalized to either GAPDH or β

### Synaptic fractionation

Synaptic plasma membrane (SPM) and postsynaptic density (PSD) fractions were prepared according to Bermejo et al. (Bermejo et al., 2014). In brief, brains of P21 mice were quickly removed and the forebrain was dissected on ice. Brain tissues were homogenized in 0.32 M sucrose in 4 mM HEPES (pH 7.4) at 900 rpm with 12 strokes using a glass-Teflon tissue homogenizer. Removal of the nuclear fraction (P1) was achieved by low speed centrifugation at 900 x g for 10 minutes. The supernatant (S1) was collected and centrifuged at 10,000 x g for 15 minutes to yield crude synaptosomal fraction (P2) and cytosol/light membranes in the supernatant (S2). P2 pellet was lysed in ddH_2_O by hypo-osmotic shock and centrifuged at 25,000 x g for 20 minutes to obtain pelleted synaptosomes (P3) and vesicular fraction (S3). The vesicular fraction was pelleted by ultracentrifugation at 165,000 x g for 2 hours. Synaptosomes (P3) were layered on top of a discontinuous sucrose gradient. Ultracentrifugation of the gradient at 150,000 x g for 2 hours yielded the SPM fraction. SPM was collected and pelleted by ultracentrifugation at 200,000 x g for 30 minutes. To prepare PSD fraction, SPM was incubated in 0.5% Triton-X100 for 15 minutes followed by centrifugation at 32,000 x g for 30 minutes. All centrifugation steps were performed at 4°C. Fractions were analyzed by Western blot and intensities of cadherin-8, cadherin-11, PSD-95 and syntaxin-1 positive signals in P3, SPM, PSD and S3 fractions were normalized to the total protein input.

### Co-immunoprecipitation

P14 forebrain and transfected and untransfected N2a cells were homogenized in lysis buffer containing 20 mM Tris, pH 7.5, 150 mM NaCl and 1% Triton X-100 supplemented with PMSF (Cell Signaling Technologies Cat#8553S) and protease and phosphatase inhibitor cocktail (Thermo Fisher Scientific Cat#78442). Protein extract (0.5 mg, standardized to 1 mg/ml) was precleared with 50 μl Protein G Sepharose 4 Fast Flow (Millipore Sigma Cat#GE17-0618-01) or μ rProtein A Sepharose Fast Flow (Millipore Sigma Cat#GE17-1279-01) for 1 hour at 4° C. g anti-cadherin-8 (DSHB Cat#CAD8-1), 7 g anti-μ cadherin-11 (Thermo Fisher Scientific Cat#32-1700) or 4 μg anti-HA antibodies (Millipore Sigma Cat#H6908), and mouse or rabbit IgGs for 2 hours at 4° C. Samples were precipitated with 50 μl of pre-equilibrated Protein G Sepharose 4 Fast Flow or rProtein A Sepharose Fast Flow for 1 hour at 4°C with gentle mixing. Immunoprecipitates were washed three times in lysis buffer and eluted by boiling in 50 μl sample buffer. Overexpressed myc-flag-tagged cadherin-8 was immunoprecipitated using the Pierce c-My-tag IP/Co-IP kit according to the manufacturer’s protocol (Thermo Fisher Cat#23620). Co-immunoprecipitated proteins were determined by Western blot analysis.

### Immunocytochemistry

Surface staining was performed on low-density hippocampal cultures (20,000 cells/2 cm^2^) at 15 DIV. Cells were washed with artificial cerebrospinal fluid (aCSF) containing 124 mM NaCl, 5 mM KCl, 1.23 mM NaH_2_PO_4_, 26 mM NaHCO_3_, 10 mM Dextrose, 1 mM MgCl_2_, 1 mM CaCl_2_, and supplemented with 10% BSA. Cells were incubated with aCSF containing primary antibody for one hour at 20°C. After washing with aCSF/10% BSA, cells were incubated with secondary antibody for one hour at 20°C before fixation with 4% paraformaldehyde for 15 minutes (Noel et al., 1999; Gu and Huganir, 2016). For total staining, N2a and primary cells were fixed with 4% paraformaldehyde, permeabilized with 0.1% Triton-X100 and incubated with primary antibodies overnight and secondary antibodies for one hour at room temperature. Cells were labeled with DAPI and coverslips were mounted on glass slides with ProLong Diamond antifade mounting solution (Thermo Fisher Scientific Cat#P36934) prior to imaging. Images were taken using a Zeiss LSM-780 scanning confocal microscope with a 40x objective/1.30 EC-plan-neofluar oil or a 63x objective/1.40 plan-apochromat oil for N2a cells, and a 63x objective/1.40 plan-apochromat oil and 4x zoom for primary cells. For co-localization analysis, cadherin-8-positive puncta were manually counted and classified as either co-localizing with PSD-95 or GAT-1 (partially or totally overlapping puncta), being adjacent to PSD-95 or GAT-1 (puncta that were in close proximity and touching each other) or cadherin-8 puncta that were PSD-95-or GAT-1-negative.

### Direct stochastic optical reconstruction microscopy (dSTORM) imaging

67,000 cells prepared from P0 C57BL/6 mouse hippocampus were plated onto Fluorodish 35 mm dishes with 23 mm #1.5-glass bottoms (Cat# FD35-100). At 15 DIV, cells were surface stained with either cadherin-8 or neuroligin-1 primary antibody and Alexa647-conjugated secondary antibody as described above. After surface staining, cells were fixed, permeabilized and immunostained with PSD-95 primary antibody and Alexa488-conjugated secondary antibody and maintained in PBS prior to imaging. Imaging was performed using a Nikon N-STORM microscope, using the following buffer system: 150 mM tris-HCl pH 8.0, 100 mM MEA-HCl (Millipore Sigma Cat#M6500), 3% Oxyfluor (Millipore Sigma Cat#SAE0059), and 2% DL-Lactate (Millipore Sigma Cat#L1375) (Nahidiazar et al., 2016). A minimum of 20,000 frames were obtained using 100% laser power of 488 and 647 nm channels with 16 ms exposure time using elliptical lens for 3D analysis. STORM processing was performed using the Fiji (NIH) plug-in ThunderSTORM (Ovesný et al., 2014). For nearest neighbor analysis, the ThunderSTORM CBC function was used to compare the cadherin-8 or neuroligin-1 localization tables to the corresponding PSD-95 table. Individual molecules of either cadherin-8 or neuroligin-1 were manually identified, and the closest molecule of PSD-95 was quantified. 50 fluorescence particles per image were examined, unless fewer than that were available for analysis. For a negative control, numbers between 0-400 were randomly generated using Microsoft Excel. For comparison between random, cadherin-8, and neuroligin-1, values were binned at 10 nm intervals and plotted using Graph Pad Prism 8 software (GraphPad Prism Software, RRID: SCR_002798).

### Morphometric analysis

Cultured *Cdh11* wild-type and knockout hippocampal neurons (300,000 cells/2 cm^2^) were transfected at 3 DIV or 9 DIV with the pLL3.7-GFP vector (Addgene Cat#11795) using Lipofectamine 2000 (Invitrogen Cat#11668-019) and fixed at 4 DIV, 7 DIV or 15 DIV with 4% paraformaldehyde, respectively. Z-stack images were taken using Zeiss LSM-780 scanning confocal microscope with a 63x objective/1.40 plan-apochromat oil at 1500x1500 resolution and stitched with 2x2 frames. For dendritic spine analysis of 15 DIV cultures, two to three dendrite segments per neuron were randomly selected from primary, secondary and tertiary branches on ≤ m in length) was quantified using Fiji (NIH). To quantify spine size, high resolution confocal images were taken with an image pixel size of 64 nm. Only secondary dendrites were analyzed. Spine size was measured using Fiji (NIH), by drawing an ROI around the entire spine, starting at the base of the spine at the dendritic shaft. For analysis of dendritic morphology, neurons were traced and the total dendrite length and branch tip number were quantified using Fiji (NIH) with NeuronJ plugin (Meijering et al., 2004). 15 DIV neurons were further analyzed using Fiji (NIH) with Sholl analysis plugin (Ferreira et al., 2014). From the center of the cell body, concentric circles having 10 µm increments in radius were defined and the number of traced dendrites crossing each circle was quantified. The complexity of dendritic arbors was analyzed by the area under the curve (AUC) using Graph Pad Prism 8 software (GraphPad Prism Software, RRID: SCR_002798).

### Calcium imaging

*Cdh11* wild-type and knockout hippocampal neurons were seeded in triplicates at a density of 30,000 cells per well on a 96-well plate coated with 20 μg/ml poly-D-lysine (Millipore Sigma Cat#P6407). At 7 DIV neurons were infected with the IncuCyte NeuroBurst Orange lentivirus under a synapsin promoter (Essen Bioscience Sartorius Cat# 4736). After 24 hours, virus was removed by changing the media. Cells were imaged at 15 DIV for 24 hours using the IncuCyte S3 system (Essen Bioscience Sartorius Cat#4763). Using the IncuCyte S3 2019A software (Essen Bioscience Sartorius) the following parameters were calculated: number of active neurons, mean correlation of activity, mean burst strength, and mean burst rate. The total number of active neurons was identified by the analysis definition. For the mean correlation of activity the temporal pattern of the change in fluorescent intensity for each active neuron was compared to every other active neuron in the image. A value between -1 and 1 was generated, with 0 being completely random (no correlation) and 1 being identical patterns of change in fluorescent intensity (highly correlated). Fisher r-to-z transformation was applied to assess the significance between correlation coefficients. The mean burst strength was analyzed by integrating the area under the curve divided by its duration. This value was calculated for each burst individually and then averaged for each active neuron, followed by averaging across the entire image. To calculate the mean burst rate the total number of bursts for each active neuron was divided by minutes of scan time, followed by averaging the values for all active neurons across the entire image. The total number of cells was obtained by counting DAPI-positive nuclei. IncuCyte NeuroLight Orange lentivirus (Essen Bioscience Sartorius Cat# 4758) was used to measure the infection rate of *Cdh11* wild-type and knockout hippocampal neurons. Neurons were infected at 7 DIV and the virus was removed after 24 hours by changing the media. Neurons were fixed with 4% paraformaldehyde at 16 DIV and imaged using Evos Auto 2.0 system (Invitrogen) with a 10x objective. The number of infected cells was counted and subsequently normalized to the total cell number determined by DAPI.

### Electrophysiology

Electrophysiological recordings were conducted using *ex vivo* brain slices of *Cdh11* wild-type and knockout littermate mice between P7 and P9 or P21 and P24. Mice were anesthetized with isoflurane, decapitated, and brains removed into ice-cold sucrose cutting solution containing (in mM): 215 Sucrose, 2.5 KCl, 1.25 NaH_2_PO4, 2.8 NaHCO_3_, 7 Dextrose, 3 Na-Pyruvate, 1 Na-Ascorbate, 0.5 CaCl_2_, 7 MgCl_2_ (pH 7.4, bubbled with 95% CO_2_/5% O_2_). Near-horizontal slices containing hippocampus, 300 µm thick, were sectioned with a vibrating microtome (VT1200S, Leica Biosystems). After cutting, slices were transferred into a warmed recovery chamber filled with bubbled artificial cerebrospinal fluid (aCSF) containing the following (in mM): 125 NaCl, 2.5 KCl, 1.25 NaH_2_PO4, 25 NaHCO_3_, 25 Dextrose, 2 CaCl_2_, 1 MgCl_2_, 3 Na-Pyruvate, and 1 Na-Ascorbate. After recovering for 30 minutes at a temperature of 34°C, the slices recovered for at least another 30 minutes at room temperature prior to recording. For recording, slices were transferred to a recording chamber superfused with the same aCSF used in the recovery chamber, maintained at 32°C. Miniature excitatory postsynaptic currents (mEPSCs) were recorded in voltage-clamp mode (V_h_ = -76 mV) in the presence of 100 µM DL-AP5, 10 µM tetrodotoxin, and 20 µM gabazine (channel blockers acquired from Tocris). Glass pipettes pulled to a resistance of 2-6 MΩ were filled with internal solution containing the following (in mM): 130 K-gluconate, 10 KCl, 10 HEPES, 0.2 EGTA, 4 MgATP, 0.5 Na_2_GTP, 10 Na_2_-phosphocreatine, and 5 QX-314 chloride, pH adjusted to 7.3 with KOH, 280–290 mOsm. Pyramidal neurons in stratum pyramidal of CA1 in dorsal hippocampal slices were visually identified for recording.

Only cells with a series resistance ≤ 30 MΩ, input resistance > 80 MΩ, and a resting membrane potential ≤ -50 mV were accepted for final analysis. Whole-cell parameters were monitored throughout the recording with a 100 ms, -10 mV step delivered every 30 sec. Recordings were made using an Axon MultiClamp 700B amplifier (Molecular Devices). Data were filtered at 2 kHz and digitized at 10 kHz with a National Instruments digital-analog converter under the control of Igor Pro software (WaveMetrics, RRID: SCR_000325). The frequency, amplitude, rise and decay times, and charge of mEPSCs were analyzed with Mini Analysis software (Synaptosoft, RRID: SCR_002184), with a threshold of 3× RM(S noise for event discrimination. In P21-P24 mice, 100-200 well-isolated events were used for these analyses in wild-type neurons; because of extremely low event frequency, a minimum of 50 events were used in knockout neurons. In P7-P9 mice, because of extremely low event frequency in CA1, all events within a 5-minute window of recording for each cell were selected for analysis.

### Ascertainment and clinical criteria of autistic and control individuals

All of the ASD patients were recruited through the University of Miami John P. Hussman Institute for Human Genomics with informed consent obtained from all participants under a University of Miami Miller School of Medicine Institutional Review Board approved protocol and all experiments were performed in compliance with the guidelines and regulations of the institutional biosafety committee. ASD individuals contributing samples for the generation of induced pluripotent stem cells (iPSCs; 110, 134, 691, 700, 709, 710, 725, 732) were ascertained following an ASD diagnosis. The core inclusion criteria were as follows: (1) individuals were between 3 and 21 years of age, (2) individuals had a presumptive clinical diagnosis of ASD, (3) an expert clinical determination of an ASD diagnosis was determined using DSM-IV (American Psychiatric Association, 2013) criteria supported by the Autism Diagnostic Interview-Revised (ADI-R; Lord, Rutter and Le Couteur, 1994), and (4) individuals had an intelligence quotient (IQ) equivalent >35 or developmental level >18 months as determined by the Vineland Adaptive Behavior Scale (VABS; Sparrow, Cicchetti and Balla, 2005). All diagnoses were based on review by a panel consisting of experienced clinical psychologists and a pediatric medical geneticist. As appropriate, IQ measures were obtained for the individuals following administration of any of several measures (e.g. age appropriate Wechsler scale, Leiter intelligence test, or Mullen Scales of Early Learning (MSEL)) or derived from medical records. All of the ASD individuals were non-syndromic and sporadic with polygenetic background as shown by whole exome sequencing (Cukier et al., 2014). The ASD lines - 110, 134, 691, 709, 725, 732 - had been derived and validated in our previous publication (DeRosa et al., 2018). Briefly, the iPSC lines were derived from peripheral blood mononuclear cells (PBMCs) isolated from the whole blood by density centrifugation. The cells were reprogrammed using the CytoTune iPS 2.0 Sendai Reprogramming Kit (Thermo Fisher Scientific Cat#A16517) according to the manufacturer’s protocol. Individual clonal iPSC lines from each participant were validated for pluripotency by immunocytochemistry for pluripotency markers (OCT3/4, NANOG, SOX2, TRA-1-81) and genomic stability confirmed by G band karyotype (WiCell) as previously published (DeRosa et al., 2018). Patients 700 and 710 were ascertained using the same approach as the other individuals and the iPSC lines derived and validated in the same manner as the previously published samples (DeRosa et al., 2018). Control samples were obtained following informed consent under an IRB approved protocol (University of Miami) from cognitively normal individuals that were between 18 and 30 years of age. These individuals had no history of ASD or other neurological disorders (e.g. schizophrenia, major depressive disorder). The control samples were slightly older than the individuals with ASD to ensure that they did not have a diagnosis of autism or a related neurodevelopmental disorder, such as schizophrenia that is often diagnosed during adolescence. PBMCs were isolated from whole blood obtained from the study participant. The reprogramming for the control lines was performed in the same manner as that used for the ASD samples. These samples were screened for pluripotency and genomic stability in the same way as the ASD samples according to our published protocol (DeRosa et al., 2018). All the samples - both the ASD and the control samples - were from non-Hispanic white (NHW) males.

### Human iPSC-derived cortical neural precursor cell cultures

iPSC lines were derived from peripheral blood obtained from eight individuals with autism and from four typically developing controls (Table 3; DeRosa et al., 2018). PBMCs were isolated and cultured in suspension until transduction using Oct4, Sox2, Klf4, and c-Myc Cytotune Sendai viruses (Thermo Fisher Scientific Cat#A16517) (DeRosa et al., 2012). After PBMC transduction into iPSCs the media was supplemented with 10 µM CHIR99021 (Stemgent Cat#04000410), 1 µM PD325901 (Stemgent Cat#040006), 1 µM thiazovivin (Stemgent Cat#040017), and 10 µM Y27632 (Stemgent Cat#04001210) for 7 days. At day 7, the media with small molecules was transitioned to mTeSR1 full stem cell maintenance media (Stemcell Technologies Cat#85850) with the media being changed daily. iPSC colonies were plated onto mouse embryonic feeders (MEFs) and grown for 7 days. Colonies were selected showing proliferating cell clusters, indicative of reprogrammed cells. To derive cortical progenitor neurons, selected iPSC colonies were dissociated via a 7-minute treatment with Accutase in the presence of 20 µM Y27632 and MEF feeders were removed using 0.1% gelatin as previously described (Nestor et al., 2015; Phillips et al., 2017). Dissociated iPSCs were exposed to media

**Table 3.** Case information of control and autism-derived iPSC lines. Sample identity number, gender, diagnosis and affected genes are listed for each iPSC line analyzed. Data was obtained from the University of Miami John P. Hussman Institute for Human Genomics (Hussman et al., 2011; Cukier et al., 2014).

|  | Sample ID | Gender | Diagnosis | Genes |
| --- | --- | --- | --- | --- |
| Autism | 110 | M | Autism | VPS13B, EFCAB5, TRIM55 |
|  | 700 | M | Autism | RBFOX1 |
|  | 725 | M | Autism | CEP290, NINL, SOS2, TRIM55, ZMYND17, BTN2A2, MDC1, FBXO40, KIAA1949, SLC8A3, TSPYL5 |
|  | 732 | M | Autism | CLCN2, F13A1, JARID2, STXBP5, C12orf73, C20orf118, FGD6 |
|  | 134 | M | Autism | CPZ, PRICKLE1, TOPOR5 |
|  | 691 | M | Autism | COL6A3, SLIT3, C2orf85, AB13BP, UIMC1 |
|  | 709 | M | Autism (Twin) | RBFOX1 |
|  | 710 | M | Autism (Twin) | RBFOX1 |
| Control | 574 | M | - | - |
|  | 321 | M | - | - |
|  | 322 | M | - | - |
|  | 324 | M | - | - |

M Y27632, 10 M SB431542 (Stemgent Cat#04001010), 1 μM thiazovivin) within growth media as described previously (DeRosa et al., 2012). After patterning and neural induction, iPSC-derived neuron progenitors were expanded using 6-well plates coated with 15 μg/ml Poly-L-Ornithine (Millipore Sigma Cat# P4957) and 10 μg/ml laminin (Invitrogen Cat# 230171015) within an enriched medium containing 1:1 mixture of DMEM/F12 (with L-Glutamine; Thermo Fisher Cat# 11320033) and Neurobasal medium (minus phenol red; Invitrogen Cat#12348017), 5μM forskolin (Millipore Sigma Cat# F6886), 60 ng/ml progesterone (Millipore Sigma Cat#P8783), 16 µg/ml putrescine (Millipore Sigma Cat#P7505), 5 μg/ml N-acetyl-L-cysteine (Millipore Sigma Cat#A8199), 1% Insulin-Transferrin-Selenium-A (Gibco Cat#41400045), 1% B-27 supplement (Gibco Cat#12587010), 0.5% N2 supplement (Gibco Cat#17502048), 1% penicillin/streptomycin (Thermo Fisher Cat#15140122), 30 ng/ml tri-iodothyronine (Millipore Sigma Cat#T6397), 40 ng/ml thyroxine (Millipore Sigma Cat#T1775), 0.5% non-essential amino acids, 100 µg/ml bovine-serum albumin (Millipore Sigma Cat#A4161) and 0.5% GlutaMAX (Invitrogen Cat#35050061). Cells were harvested at 19 DIV in RIPA buffer (Cell Signaling Technologies Cat# 9806S) supplemented with PMSF (Cell Signaling Technologies Cat#8553S) and protease and phosphatase inhibitor cocktail (Thermo Fisher Scientific Cat#78442) and triplicates of each control and autism line were analyzed by Western blot.

### Organoid culture and mRNA quantitation via qPCR

Cortical organoid cultures were generated from four control and six autism lines (Table 3) as previously described in Durens et al. (Durens et al., 2020). In brief, PBMC-derived iPSCs were grown on MEF feeder layers and neural induction was performed as described above. At 14 DIV, aggregates were transferred to 0.4 μm PTFE inserts (Millipore Cat#PICM0RG50) and cultured in enriched medium containing DMEM/F12 medium (Gibco Cat# 11320033) and Neurobasal medium (1:1; Gibco Cat#12348017), 2% B27 supplement (Gibco Cat#12587010), 0.5% N2 supplement (Gibco Cat#17502048), 0.5% non-essential amino acids (Gibco Cat#11140050), 1% SATO mix (progesterone (Millipore Sigma Cat#P8783), putrescine (Millipore Sigma Cat#P7505), tri-idothyronine (Millipore Sigma Cat#T6397), thyroxine (Millipore Sigma Cat#T1775), and bovine-serum albumin (Millipore Sigma Cat#A4161), N-acetyl-L-cysteine (Millipore Sigma Cat#A8199), 1% Insulin-Transferrin-Selenium-A (Thermo-Fisher Cat#51300044), 1% penicillin/streptomycin (Thermo Fisher Cat#15140122), and supplemented with the following neurotrophic factors: 20ng/ml BDNF (Peprotech Cat#45002), 20ng/ml NT3 (Peprotech Cat#45003), 20ng/ml β μg/ml laminin (ThermoFisher Cat#23017015), 5 M forskolin (Millipore Sigma Cat#F6886), and 2 μg/ml heparin (Millipore Sigma Cat#H3149). At 30 DIV, 2-NGF and heparin were removed to induce terminal differentiation. At 45 DIV, DAPT was removed. Organoids were cultured for a total of 60 days, and then transferred to RNAprotect Cell Reagent (Qiagen Cat#76526) for RNA stabilization. Two organoids were pooled per biological replicate and triplicates of each control and autism line were analyzed. Samples were stored at -80°C until processed. Samples in RNAprotect were thawed on ice and centrifuged at 5000 x g for 5 minutes after which supernatant was removed. Total RNA was extracted using the RiboPure Kit (Thermo Fisher Scientific Cat#AM1924) following manufacturer’s specifications. RNA concentration was determined using NanoDrop 8000 Microvolume UV-Vis Spectrophotometer (Thermo Fisher Scientific). DNA-*free* DNA removal kit (Thermo Fisher Scientific Cat#AM1906) was used to remove residual DNA. Reverse transcription (RT) was performed using 250 ng of DNase-treated RNA using the iScript cDNA synthesis kit (Bio-Rad Cat#1708890) and the resulting RT reactions were diluted 1:5 with nuclease-free water. qPCR was performed for the following genes: CDH8 (Forward – 5’-ACA GCG AAT TTT GAA CCG CTC-3’; Reverse – 5’-TCC TCC CGG TCA AGT CTT TTT -3’) and CDH11 (Forward – 5’-AGA GGT CCA ATG TGG GAA CG -3’, Reverse: 5’-GGT TGT CCT TCG AGG ATA CTG -3’). GAPDH was used as a reference gene (Genecopoeia Catalog#: HQP006940). qPCR was performed with 2X All-in-One qPCR Mix (Genecopoeia Catalog#: QP001) using the following reaction mix: 10µl All-in-One qPCR Mix (2x), 2 µl of 20µM primer, 5µl nuclease-free water, and 5 µl of cDNA at 1:5 dilution. Reactions were incubated at 95° for 10 minutes, followed by 40 cycles of 95° for 10 seconds, 60° for 20 seconds, and 72° for 30 seconds using the CFX96 Real-time System (Bio Rad). Melt curves were generated after amplification by increasing temperature from 72° to 95° at 0.5° increments. All reactions were performed in duplicates. Relative quantities for each gene were analyzed using the comparative Ct method. Fold changes were calculated relative to the average Ct values for control samples.

### Experimental design and statistical analysis

All data are reported as mean ± standard error of the mean (SEM) as either line graphs or bar graphs with scatter plots, or as box and whisker plots displaying all data points from minimum to maximum. Statistical analysis was performed using Graph Pad Prism 8 software (GraphPad Prism Software, RRID: SCR_002798). Unpaired two-tailed *t*-test was performed when comparing two groups and one-way ANOVA with Tukey’s or Dunnett’s multiple comparisons test was used to compare differences between three to six groups. *P-*values were considered significant if ≤ 0.05. All *p-*values, t-values, F-values, degrees of freedom (DF), sample sizes (N) as well as statistical tests performed are reported in Table 5 and in the figure legends. Sample sizes were predetermined before each experiment. Normality tests were run to confirm normal distribution of the samples.

## Results

### Cadherin-8 and cadherin-11 show similar temporal and spatial expression patterns

We first compared the overall protein levels of classical type II cadherins, cadherin-8 and cadherin-11, in C57BL/6 wild-type mouse whole brain samples taken at different ages of development from embryonic day (E) 14 to postnatal day (P) 21, as well as in adulthood, by Western blot (Figure 1A). The anti-cadherin-8 and anti-cadherin-11 antibodies were validated for specificity by Western blot (Figure 1-1A, B). Cadherin-8 and cadherin-11 both exhibited relatively low expression at E14 but their levels increased postnatally (Figure 1B). Cadherin-8 levels readily increased at P1 and reached more than a 10-fold increase by P7 and P14. The expression of Cadherin-8 dropped after P14 and remained at low levels in adulthood. Cadherin-11 levels reached an approximately 2-fold increase by P7 and P14 compared to E14, but were reduced to P1 level by P21 and to E14 level by adulthood (Figure 1B). Comparison to the developmental timeline showed that cadherin-8 and cadherin-11 expression correlated to time points during which processes of dendrite development and synaptogenesis are prevalent, suggesting that they may play a role in these processes. We then analyzed the protein levels in specific brain regions at P14 (Figure 1C). Cadherin-8 expression was significantly higher in the cortex, hippocampus and thalamus/striatum compared to the cerebellum (Figure 1D). Cadherin-11 showed a similar expression pattern, although differences between regions were not statistically significant (Figure 1E).

To further interrogate which brain cell type(s) express cadherins, protein levels of cadherin-8 and cadherin-11 were measured in primary cortical neurons and non-neuronal cells cultured from mouse cortices. The temporal expression patterns of these two cadherins in primary cortical neurons showed a gradual and significant increase of cadherin-8 and a trend towards increased expression of cadherin-11 from 1 DIV to 14 DIV (Figure 2A, B). In contrast to neurons, cadherin-8 was virtually undetectable in primary non-neuronal cells (Figure 2C). Cadherin-11 was expressed in both cortical neurons and non-neuronal cells, but the protein levels in neurons were two-fold greater than in non-neuronal cells (Figure 2D). These results demonstrate that both cadherins are preferentially expressed in neurons.

**Figure 2.**
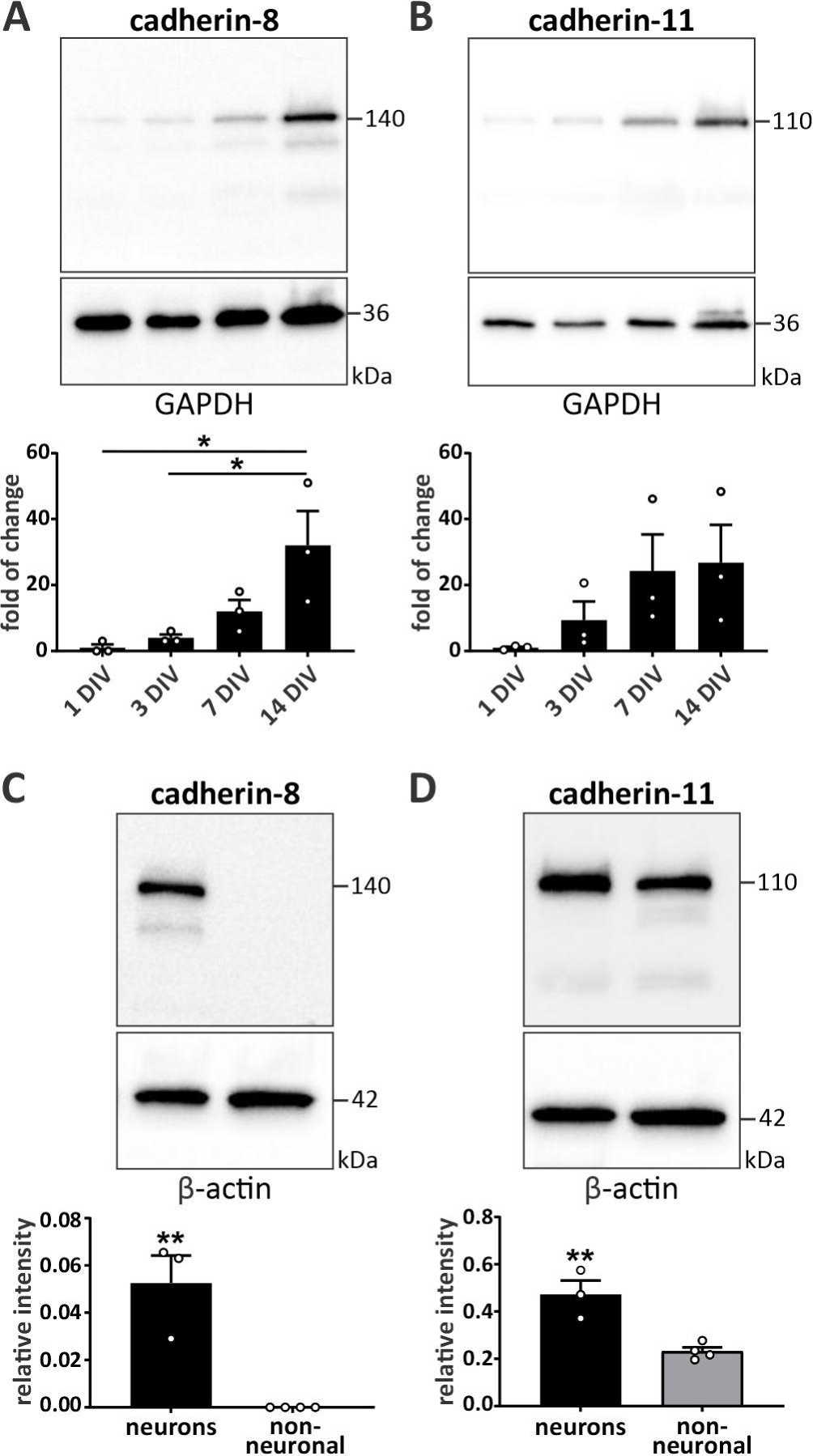
Cadherin-8 and cadherin-11 are preferentially expressed in neurons. Temporal expression pattern of **(A)** cadherin-8 and **(B)** cadherin-11 in cortical neurons harvested at 1, 3, 7 and 14 DIV. \**p* = 0.0177 between 1 DIV and 14 DIV, \**p* = 0.0299 between 3 DIV and 14 DIV, one-way ANOVA with Tukey’s multiple comparisons test. N = 3 independent cultures. Relative expression levels of **(C)** cadherin-8 and **(D)** cadherin-11 in cultured neurons and non-neuronal cells. \*\**p* < 0.01, unpaired two-tailed *t*-test. N = 3 neuronal and 4 non-neuronal cell cultures. Cells were cultured for 14 DIV before harvest. Cadherin signals were normalized to GAPDH (A,B) and β-actin (C, D).

### Cadherin-8 and cadherin-11 localize to synaptic compartments but reveal distinct interaction profiles

We next performed subcellular fractionation of forebrain tissue to isolate synaptic plasma membrane (SPM) and postsynaptic density (PSD) to determine whether cadherins are enriched in synaptic compartments (Figure 3A, B; Table 4). The distribution of PSD-95, syntaxin-1, and -β actin were evaluated to confirm successful separation and purity of different subcellular fractions. PSD-95 was used as a positive control for protein enrichment in SPM and PSD fractions - it was significantly enriched in SPM and PSD fractions compared to total protein input (Figure 3B; Table 4). Syntaxin-1 was significantly enriched in SPM and, consistent with its presynaptic expression and function in vesicle release, expression in synaptic vesicle fraction (S3) was higher compared to PSD fraction (Figure 3B; Table 4). Cadherin-8 showed a trend of enrichment in SPM and PSD, although the difference was not significant (Figure 3B; Table 4). Cadherin-11 showed significant enrichment in both, SPM and PSD fractions (Figure 3B; Table 4). Together, these results show that both cadherin-8 and cadherin-11 are present in pre- as well as postsynaptic membranes and localize to postsynaptic densities.

**Figure 3.**
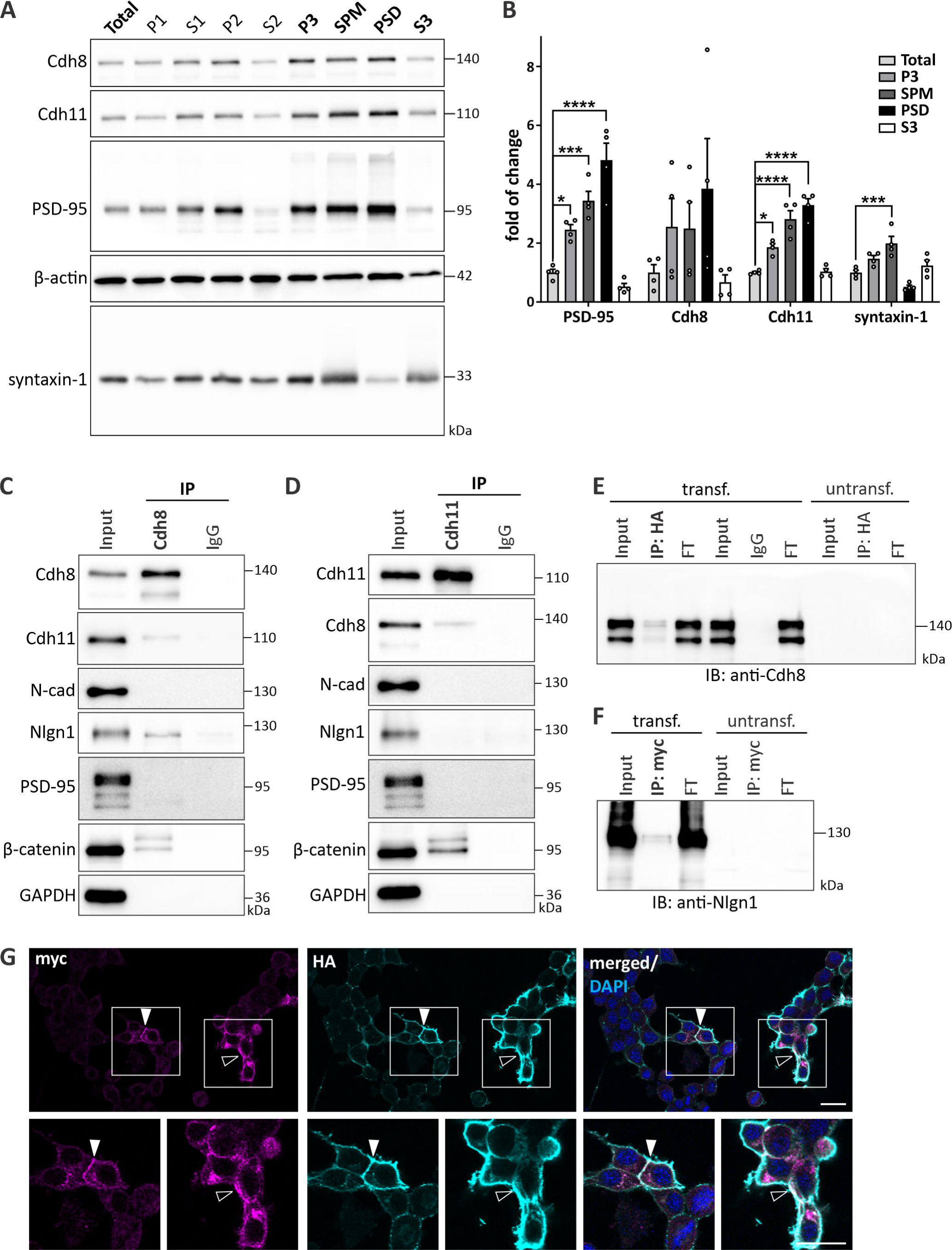
Cadherin-8 and cadherin-11 are expressed in synaptic compartments and cadherin-8 interacts with neuroligin-1. **(A)** Forebrain tissues were subjected to synaptic fractionation analysis to determine the subcellular localization of cadherin-8 and cadherin-11. Markers (PSD-95 and syntaxin-1) were probed as control for purity of the fractionation. Total: total protein input, P1: nuclear, S1: cytosol/membranes, P2: crude synaptosome, S2: cytosol/light membranes, P3: synaptosome, SPM: synaptic plasma membrane, PSD: postsynaptic density, S3: synaptic vesicles. **(B)** Quantification of protein enrichment in P3, SPM, PSD and S3 fractions compared to total protein input. PSD-95: \**p* = 0.0159, \*\*\**p* = 0.0002, \*\*\*\**p* < 0.0001; Cdh8: *p* = 0.6618 (Total vs. SPM), *p* = 0.1673 (Total vs. PSD); Cdh11: \**p* = 0.0113, \*\*\*\**p* < 0.0001; Syntaxin-1: \*\*\**p* = 0.0009; one-way ANOVA with Dunnett’s multiple comparisons test. N = 8 P21 mice, two pooled forebrains per sample. Representative Western blot of immunoprecipitation (IP) of **(C)** cadherin-8 and **(D)** cadherin-11 from P14 forebrain tissues. Cadherin-8 binds to cadherin-11, β-catenin and neuroligin-1 whereas cadherin-11 binds to cadherin-8 and -catenin, but not neuroligin-1. N = 3 independent co-IPs. Representative Western blot of **(E)** IP of neuroligin-1-HA immunoblotted for Cdh8 and **(F)** IP of cadherin-8-myc immunoblotted for Nlgn1 from N2a cells over-expressing cadherin-8-myc and neuroligin-1-HA. Cadherin-8-myc is present in the IP of neuroligin-1-HA and neuroligin-1-HA is present in the IP of cadherin-8-myc. Cadherin-8 and neuroligin-1 are not expressed endogenously in N2a cells. FT **=** flow-through. N = 3 independent co-IPs. **(G)** Immunocytochemistry of N2a cells over-expressing cadherin-8-myc (magenta) and neuroligin-1-HA (cyan), counterstained for DAPI (blue). Cadherin-8-myc and neuroligin-1-HA show partial co-localization at cell-cell contacts (arrowheads) and at the cell membrane (open arrowheads). Boxed areas are magnified.Scale bar = 20 μm.

**Table 4.**
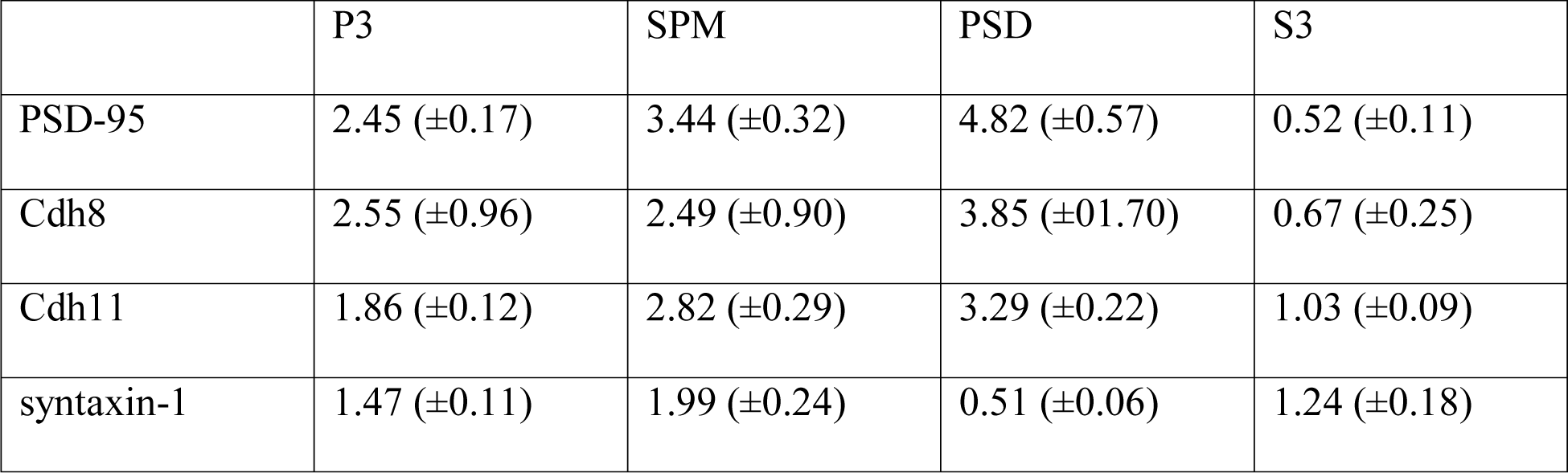
Enrichment of protein expression in synaptic fractions. Mean values (±SEM) of enrichment of PSD-95, cadherin-8, cadherin-11 and syntaxin-1 in different subcellular fractions, including synaptosome (P3), synaptic plasma membrane (SPM), postsynaptic density (PSD) and synaptic vesicle fraction (S3).

**Table 5.** Statistical results. Detailed information about statistical test, F-values, t-values, p-values and degree of freedom (DF) for all experiments performed in this study.

| Figure | Test | F (DFn, DFd) | t-value | *p*-value | DF |
| --- | --- | --- | --- | --- | --- |
| 1B, Cdh8 | One-way ANOVA | F (5, 30) = 10.26 |  | <0.0001 | 5 |
| 1B, Cdh11 | One-way ANOVA | F (5, 30) = 3.384 |  | 0.0153 | 5 |
| 1D | One-way ANOVA | F (3, 12) = 7.040 |  | 0.0055 | 3 |
| 1E | One-way ANOVA | F (3, 12) = 2.176 |  | 0.1438 | 3 |
| 2A | One-way ANOVA | F (3, 8) = 6.339 |  | 0.0165 | 3 |
| 2B | One-way ANOVA | F (3, 8) = 2.104 |  | 0.1781 | 3 |
| 2C | *t*-test |  | 5.332 | 0.0031 | 5 |
| 2D | *t*-test |  | 4.534 | 0.0062 | 5 |
| 3B, PSD-95 | One-way ANOVA | F (4, 15) = 32.22 |  | <0.0001 | 4 |
| 3B, Cdh8 | One-way ANOVA | F (4, 15) = 1.747 |  | 0.1921 | 4 |
| 3B, Cdh11 | One-way ANOVA | F (4, 15) = 34.99 |  | <0.0001 | 4 |
| 3B, syntaxin-1 | One-way ANOVA | F (4, 15) = 13.81 |  | <0.0001 | 4 |
| 4C | One-way ANOVA | F (2, 114) = 10.10 |  | <0.0001 | 2 |
| 4D | One-way ANOVA | F (2, 90) = 78.09 |  | <0.0001 | 2 |
| 4J | One-way ANOVA | F (2, 57) = 36.49 |  | <0.0001 | 2 |
| 5A | *t*-test |  | 3.950 | 0.0027 | 10 |
| 5B | *t*-test |  | 2.485 | 0.0323 | 10 |
| 5C | *t*-test |  | 2.342 | 0.0473 | 8 |
| 5D | *t*-test |  | 6.748 | 0.0001 | 8 |
| 6B, primary | *t*-test |  | 1.128 | 0.2638 | 59 |
| 6B, secondary | *t*-test |  | 1.346 | 0.1833 | 59 |
| 6B, tertiary | *t*-test |  | 0.9198 | 0.3614 | 59 |
| 6B, total | *t*-test |  | 0.4928 | 0.6239 | 59 |
| 6C | *t*-test |  | 0.5255 | 0.6014 | 54 |
| 6E | *t*-test |  | 2.286 | 0.0269 | 46 |
| 6F, TDL | *t*-test |  | 2.207 | 0.0323 | 46 |
| 6F, tip number | *t*-test |  | 2.387 | 0.0212 | 46 |
| 6G, TDL | *t*-test |  | 1.745 | 0.0845 | 88 |
| 6G, tip number | *t*-test |  | 0.5062 | 0.6140 | 88 |
| 6H, TDL | *t*-test |  | 0.2949 | 0.7689 | 68 |
| 6H, tip number | *t*-test |  | 0.8764 | 0.3839 | 68 |
| 6J | *t*-test |  | 10.06 | <0.0001 | 12 |
| 6K | *t*-test |  | 2.945 | 0.0123 | 12 |
| 6L | *t*-test |  | 4.004 | 0.0017 | 12 |
| 6M | *t*-test |  | 3.496 | 0.0044 | 12 |
| 6N | *t*-test |  | 0.9961 | 0.3389 | 12 |
| 6O, PSD-95 | *t*-test |  | 0.2046 | 0.8420 | 10 |
| 6O, Nlgn1 | *t*-test |  | 0.05644 | 0.9561 | 10 |
| 6O, Cdh8 | *t*-test |  | 1.582 | 0.1447 | 10 |
| 6O, N-cadherin | *t*-test |  | 0.3710 | 0.7184 | 10 |
| 6P, PSD-95 | *t*-test |  | 1.944 | 0.0805 | 10 |
| 6P, Nlgn1 | *t*-test |  | 0.7755 | 0.4560 | 10 |
| 6P, Cdh8 | *t*-test |  | 1.674 | 0.1252 | 10 |
| 6P, N-cadherin | *t*-test |  | 0.9067 | 0.3859 | 10 |
| 7B | *t*-test |  | 3.481 | 0.0022 | 21 |
| 7C | *t*-test |  | 3.427 | 0.0027 | 20 |
| 7D | *t*-test |  | 2.541 | 0.0259 | 12 |
| 7E | *t*-test |  | 0.8066 | 0.4290 | 21 |
| 7G | *t*-test |  | 2.254 | 0.0456 | 11 |
| 7H | *t*-test |  | 2.234 | 0.0472 | 11 |
| 7I | *t*-test |  | 3.796 | 0.0030 | 11 |
| 7J | *t*-test |  | 1.289 | 0.2238 | 11 |
| 8B | *t*-test |  | 0.1577 | 0.8775 | 11 |
| 8C | *t*-test |  | 0.4992 | 0.6260 | 13 |
| 8D | *t*-test |  | 2.591 | 0.0160 | 24 |
| 8E | *t*-test |  | 3.101 | 0.0049 | 24 |
| 8F | *t*-test |  | 3.947 | 0.0006 | 24 |
| 8G | *t*-test |  | 0.7550 | 0.4576 | 24 |
| 9C | *t*-test |  | 0.5769 | 0.5688 | 27 |
| 9D | *t*-test |  | 0.3030 | 0.7642 | 27 |
| 9E | *t*-test |  | 0.4087 | 0.6860 | 27 |
| 9F | *t*-test |  | 0.5385 | 0.5946 | 27 |
| 9G | *t*-test |  | 0.2913 | 0.7731 | 27 |
| 9J | *t*-test |  | 4.776 | <0.0001 | 46 |
| 9K | *t*-test |  | 2.486 | 0.0166 | 46 |
| 9L | *t*-test |  | 1.984 | 0.0532 | 46 |
| 9M | *t*-test |  | 0.0004 | 0.9997 | 46 |
| 9N | *t*-test |  | 0.5073 | 0.6144 | 46 |

Since cadherin-8 and cadherin-11 were both expressed in the PSD fraction, we performed co-immunoprecipitation (co-IP) experiments using anti-Cdh8 (Figure 3C) and anti-Cdh11 antibodies (Figure 3D) to determine the association of these two cadherins with specific postsynaptic proteins. Both, cadherin-8 and cadherin-11, immunoprecipitated β-catenin, the conserved intracellular binding partner of classical cadherins (Seong et al., 2015). Also, both cadherins interacted with each other, consistent with previous reports showing cadherin-8-cadherin-11 interactions in cultured cells and through biophysical analysis (Shimoyama et al., 2000; Brasch et al., 2018). Neither cadherin-8 nor cadherin-11 co-immunoprecipitated with PSD-95 or N-cadherin. Previous studies found cooperative functions of classical cadherins with the excitatory synaptic adhesion molecule neuroligin-1 (Nlgn1) during neural circuit development (Stan et al., 2010; Aiga et al., 2011; Yamagata et al., 2018). This prompted us to investigate whether cadherin-8 and cadherin-11 physically interact with neuroligin-1. Interestingly, neuroligin-1 was detected in the immunoprecipitate of cadherin-8 (Figure 3C), but not cadherin-11 (Figure 3D). We confirmed this interaction *in vitro* by co-expressing myc-tagged *Cdh8* together with HA-tagged *Nlgn1* in N2a cells that do not endogenously express these proteins (Figure 3E, F, Figure 1-1C, D). Using either anti-HA or anti-myc antibodies to selectively pull down neuroligin-1 and cadherin-8, respectively, we found that neuroligin-1 co-immunoprecipitated cadherin-8 and vice versa. Consistent with these findings, immunostaining of N2a cells overexpressing myc-tagged *Cdh8* and HA-tagged *Nlgn1* showed a partial co-localization of these two proteins at the plasma membrane and at cell-cell contacts (Figure 3G).

The finding of an interaction of cadherin-8 with neuroligin-1 prompted us to further examine the subcellular localization of cadherin-8 in neurons. The specificity of the anti-cadherin-8 antibody was first validated in immunocytochemistry (Figure 1-1C).

Immunofluorescence of surface-exposed cadherin-8 in 15 DIV hippocampal neurons revealed partial co-localization with the excitatory synaptic markers PSD-95 and synapsin-1 (Figure 4A). The majority of cadherin-8 localized to excitatory synaptic puncta, either co-localizing with PSD-95 (30.3% ± 2.0%) or located directly adjacent to PSD-95-positive puncta (41.6% ± 2.5%), while the remainder of cadherin-8 puncta (28.1% ± 2.2%) did not localize to PSD-95-positive puncta (Figure 4C). Hence, most, but not all, cadherin-8-positive puncta are associated with excitatory synapses. Conversely, the majority of the cadherin-8-positive puncta (56.8% ± 3.4%) did not associate with GAT-1-positive puncta, markers of inhibitory synapses (Figure 4B and D). However, a small fraction of cadherin-8-positive puncta exhibited co-localization with (10.0% ± 1.8%) or were adjacent to GAT-1-positive puncta (33.2% ± 2.6%). These data suggest that cadherin-8 is enriched at/near excitatory synapses compared to inhibitory synapses, consistent with the potential role in excitatory synaptic function. To further quantify cadherin-8 presence at excitatory synapses we performed direct stochastic reconstruction microscopy (dSTORM). We examined the distance between PSD-95 and either cadherin-8 or neuroligin-1, a positive control for excitatory synaptic localization (Figure 4E, F). For optimal two-color imaging, we used an Oxyfluor buffer system (Nahidiazar et al., 2016). 33.3% ± 3.3% of cadherin-8 puncta were located within 50 nm of a PSD-95 puncta, significantly more than a randomized set of data points (Figure 4G, I, J; 13.7% ± 1.0%). This recapitulates the co-localization data obtained using standard confocal microscopy. Interestingly, dSTORM imaging showed that significantly more neuroligin-1 puncta were within 50 nm of a PSD-95 punctum compared to cadherin-8 puncta and the randomized data points (Figure 4H, I, J; 52% ± 3.8%). Together, these data indicate that cadherin-8 localizes to excitatory synapses but to a lesser extent than neuroligin-1. These findings are in line with the localization of cadherin-8 and neuroligin-1 in N2a cells suggesting that these two proteins partially but not completely localize to the same cellular compartment.

**Figure 4.**
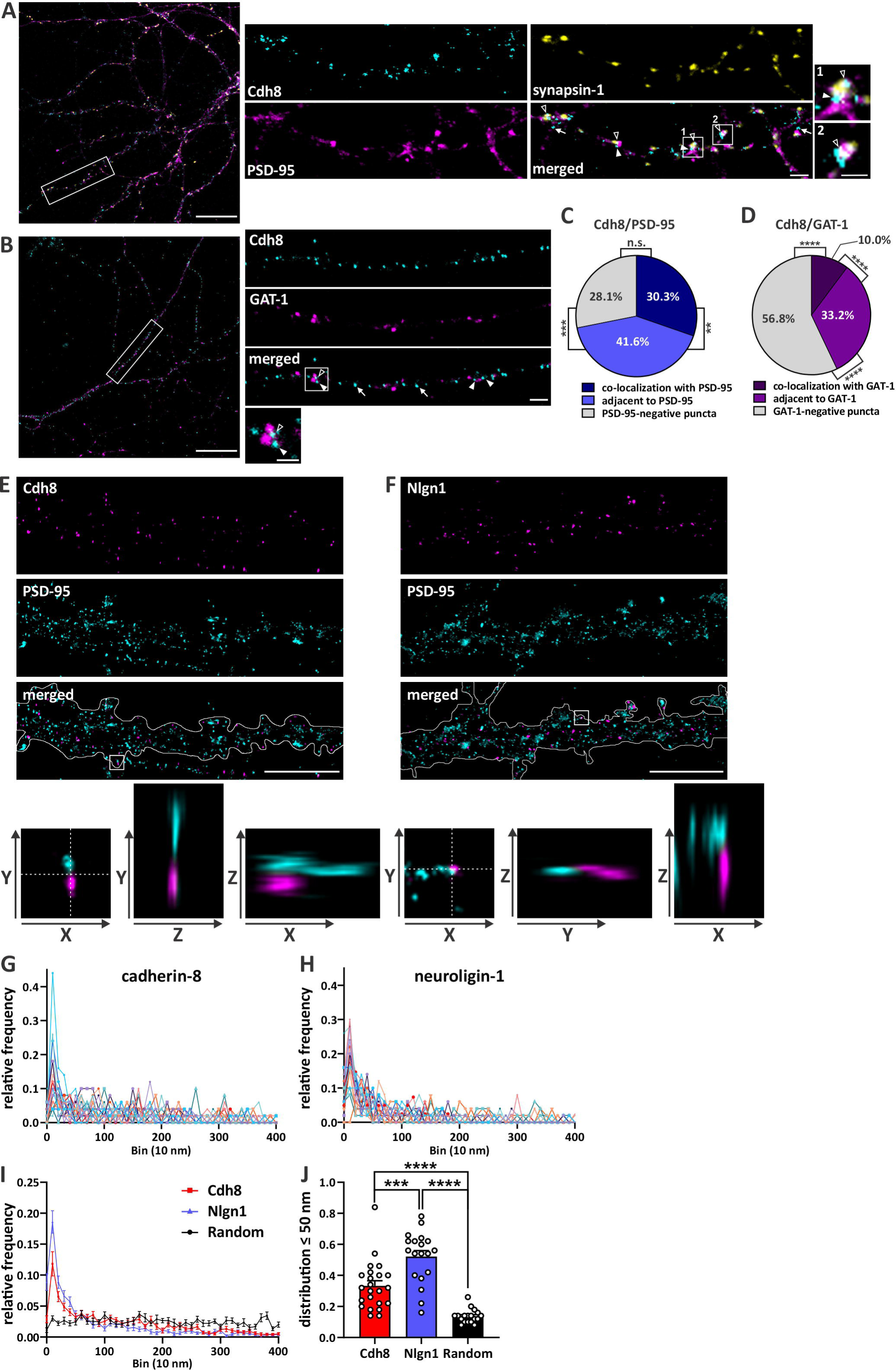
Cadherin-8 localizes to excitatory synapses. Representative confocal images showing surface expressed cadherin-8 (cyan) on dendritic segments of cultured hippocampal neurons at 15 DIV co-immunostained with either the excitatory synaptic markers **(A)** PSD-95 (magenta) and synapsin-1 (yellow), or the inhibitory synaptic marker **(B)** GAT-1 (magenta). Cadherin-8 puncta co-localizing with PSD-95-or GAT-1-positive puncta (open arrowheads), being adjacent to PSD-95-or GAT-1-positive puncta (arrowheads) and PSD-95-or GAT-1-m (overview), 2 μm (dendritic segments). Representative synaptic puncta are magnified (boxed areas, scale bar = 1 μm). The synaptic profile of cadherin-8-positive puncta were quantified and analyzed as fractions of co-localization with **(C)** PSD-95 or **(D)** GAT-1. PSD-95: *\*\*p* = 0.0019, \*\*\**p* = 0.0002, GAT-1: *p* < 0.0001, one-way ANOVA with Tukey’s multiple comparisons test. N = 11-14 neurons, three independent cultures. dSTORM imaging was performed on 15 DIV hippocampal neurons immunostained for PSD-95 (cyan) and either surface-expressed **(E)** cadherin-8 (magenta) or **(F)** neuroligin-1 (magenta). Dendrites are outlined in merged images. Scale bar = 5 µm. Insets (800 nm) show 3D proximity of PSD-95 to the surface stained proteins. Crosshair shows YZ and XZ planes. **(G-J)** Nearest neighbor analysis of dSTORM data. Individual surface stained **(G)** cadherin-8 and **(H)** neuroligin-1 puncta were isolated, and the nearest PSD-95 puncta was determined. **(I)** Quantification and comparison of nearest neighbor analysis for cadherin-8, neuroligin-1 and a randomized set of numbers generated between 0-400 nm. **(J)** Quantification and comparison of the frequency distribution of cadherin-8 and neuroligin-1 puncta and a randomized data set between 0 and 50 nm from the nearest PSD-95 puncta. \*\*\**p* = 0.0001, \*\*\*\**p* < 0.0001, one-way ANOVA with Tukey’s multiple comparisons test. N = 23 neurons (Cdh8) and 19 neurons (Nlgn1), 18 random data points, three independent cultures. 50 particles per cell were used, unless fewer particles were available.

### CDH8 and CDH11 levels are altered in autism-specific iPSC-derived cortical neuronal progenitor cells and organoids

CDH8 and CDH11 have been identified as autism-risk candidate genes (Hussman et al., 2011,; Pagnamenta et al., 2011; Cukier et al., 2014; Crepel et al., 2014). Our findings of a dynamic expression profile of both cadherin-8 and cadherin-11 in mouse brain development and their (sub)-cellular localization in neurons and to synaptic compartments indicates that these two cadherins may function in neural circuit formation. Impaired neuronal connectivity has been suggested to represent a risk factor contributing to etiology of autism (Betancur et al., 2009; Bourgeron, 2009; Hussman et al., 2011), which prompted us to investigate the involvement of cadherin-8 and cadherin-11 in autism. As autism is a genetically complex condition with broad heterogeneity across individuals, we wanted to determine whether the levels of these cadherins are commonly altered in autism. We thus investigated the expression of CDH8 and CDH11 in human tissue using samples from autistic and typically-developing control individuals. iPSC-derived cortical neural progenitor cells (NPCs) from four neurotypical control and eight autistic individuals were cultured for 19 DIV and cell lysates were harvested for protein analysis (Table 3; Figure 5A, B, Figure 1-1A, B). Although none of these individuals carried risk variants in either of these cadherins (DeRosa et al., 2018), CDH8 levels were significantly increased (Figure 5A; ctrl: 0.0614 ± 0.0108, autism: 0.1186 ± 0.0086) whereas CDH11 levels were significantly decreased (Figure 5B; ctrl: 0.4200 ± 0.1520, autism: 0.0821 ± 0.0621) in iPSC-derived cortical NPCs from autistic individuals compared to neurotypical control cells. We next performed quantitative PCR (qPCR) to analyze mRNA expression of CDH8 and CDH11 in iPSC-derived cortical organoids generated from six autistic and four typically-developing control individuals that were grown in culture for 60 DIV (Table 3; Figure 5C, D). These cortical organoids mimic early cortical development with active excitatory and inhibitory neurotransmission (Durens et al., 2020). In line with the expression profile in the NPCs, autism-specific iPSC-derived cortical organoids showed a significant increase of CDH8 (Figure 5C; ctrl: 1.072 ± 0.1783, autism: 3.093 ± 0.6812) and a concomitant decrease of CDH11 (Figure 5D; ctrl: 1.016 ± 0.0249, autism: 0.5603 ± 0.0517) compared to neurotypical control organoids. Together, these results show that both cadherin-8 and cadherin-11 levels are altered in cells derived from individuals with autism that mimic early stages of neural circuit development.

**Figure 5.**
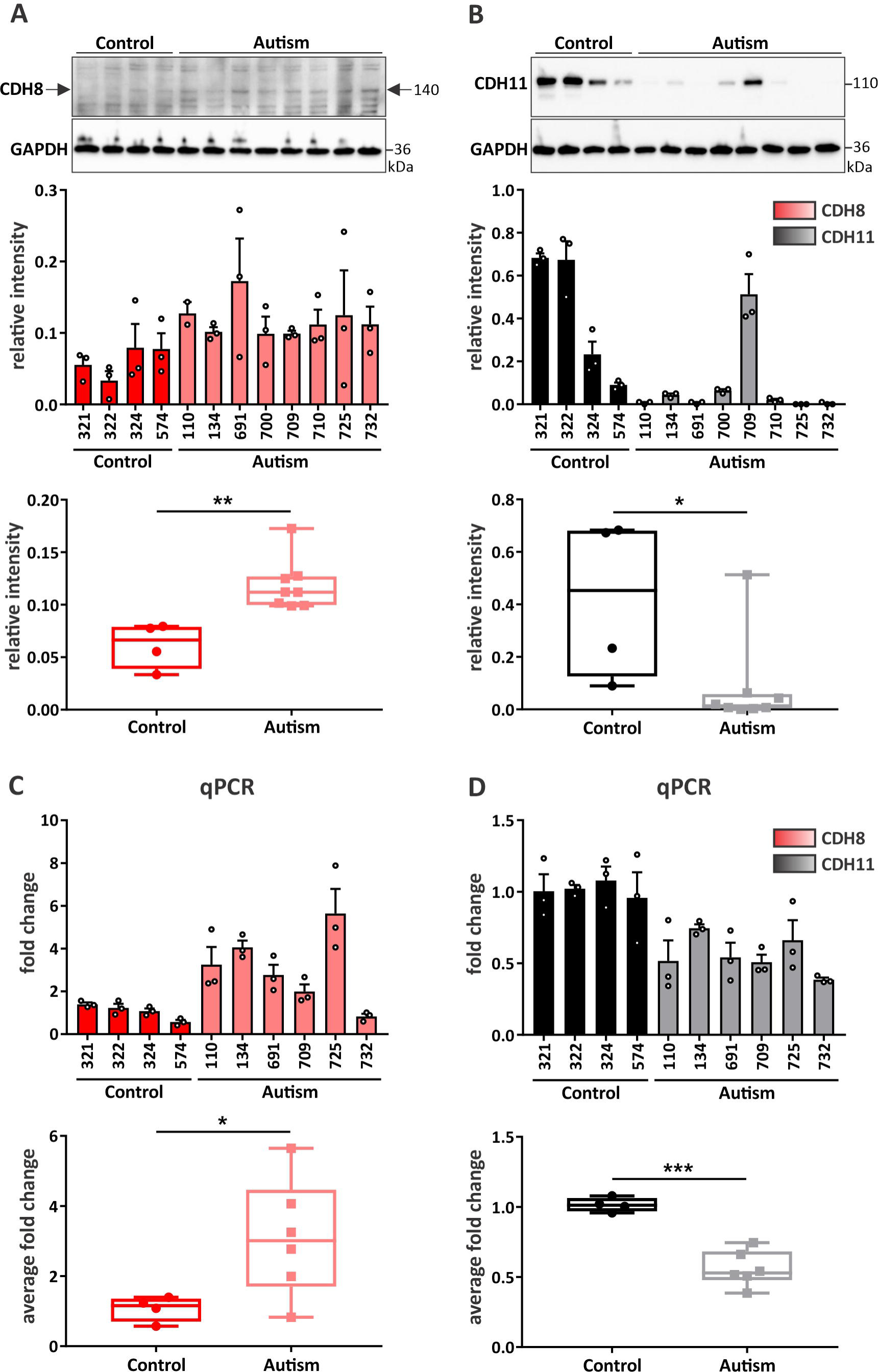
iPSC-derived cortical NPCs and organoids from autistic individuals show altered expression of CDH8 and CDH11. Western blot analysis and quantification of **(A)** CDH8 and **(B)** CDH11 expression in iPSC-derived cortical NPCs from typically-developing control and autistic individuals at 19 DIV. Quantification of the expression of each cadherin is represented as bar graph and as box and whisker plot. \**p* < 0.05, \*\**p* < 0.01; unpaired two-tailed *t*-test. N = 4 control and 8 autistic individuals, triplicates of each individual. Cadherin signals were normalized to GAPDH. Quantification of mRNA expression of **(C)** CDH8 and **(D)** CDH11 in cortical organoids derived from iPSCs of control and autistic individuals at 60 DIV via qPCR. \**p* < 0.05, \*\*\**p* < 0.001; unpaired two-tailed *t*-test. N = 4 control and 6 autism individuals, triplicates of each individual.

### Using *Cdh11^-/-^* mice as a model to study potential autism-related phenotypes show increased dendrite complexity in neurons

The *Cdh11* knockout mouse has been previously shown to exhibit behaviors that are similar to autism phenotypes (Horikawa et al., 1999; Wu et al., 2020). The significant downregulation of CDH11 in autism-specific NPCs prompted us to use the *Cdh11* knockout mice as a model system to examine potential pathological phenotypes during neural circuit development and their underlying mechanisms in autism. Alterations in excitatory synapse development is one of the hallmarks of autism (Hutsler and Zhang, 2010; Forrest et al., 2018). Since we observed a peak of cadherin-11 expression during the time frame of synaptogenesis and enrichment in PSD fractions (Figure 1B, 3B), we first examined the density of dendritic spines, the major structures that harbor excitatory synapses, in 15 DIV hippocampal cultures from *Cdh11* wild-type and knockout mice (Figure 6A). Intriguingly, we did not observe changes in the dendritic spine density on primary, secondary and tertiary dendrites of *Cdh11^-/-^* neurons compared to wild-type neurons (Figure 6B; Primary – WT: 0.7664 ± 0.0301 μm^-1^, KO: 0.7086 ± 0.0428 μm^-1^; Secondary – WT: 0.7070 ± 0.0270 μm^-1^, KO: 0.6527 ± 0.0301 μm^-1^ ; Tertiary – WT: 0.6051 ± 0.0265 μm^-1^, KO: 0.6440 ± 0.0337 μm^-1^ ; Total – WT: 0.6778 ± 0.0194 μm^-1^, KO: 0.6607 ± 0.0299 μm^-1^). We also did not find significant differences in the size of dendritic spines on secondary dendrites from *Cdh11^-/-^* neurons compared to wild-type neurons (Figure 6C; WT: 0.2970 ± 0.0123m^2^, KO: μ 0.2870 ± 0.0140 μ). We then analyzed the dendritic morphology of *Cdh11* knockout and wild-type neurons from 15 DIV hippocampal cultures (Figure 6D). Sholl analysis revealed a significant increase in the dendritic arbor complexity in *Cdh11^-/-^* neurons compared to wild-type neurons (Figure 6E; area under the curve (AUC), WT: 2,551 ± 153.1, KO: 3,051 ± 155.5). The increase in dendritic complexity resulted from a significant increase in total dendrite length as well as branches as revealed by branch tip number (Figure 6F; TDL – WT: 2,509 ± 174 μm, KO: 3,034 ± 162.7 μm; branch tip number – WT: 29.17 ± 2.194, KO: 35.4 ± 1.48). To evaluate whether the dendritic phenotype was readily present at earlier developmental time points, we further analyzed total dendrite length and branch tip number in neurons fixed at 4 DIV and 7 DIV. In contrast to 15 DIV, total dendrite length and branch tip number of *Cdh11^-/-^* neurons were not significantly increased compared to wild-type neurons at both time points (Figure 6G, μm, KO: 641.7 ± 31.51 m, branch tip number WT: 16.53 ± m, KO: 1,129 ± 81.23 μm, branch tip number WT: 21.15 ± 1.312, KO: 22.76 ± 1.274). Together, these data indicate that loss of cadherin-11 leads to altered morphology of dendrites during a later developmental period.

**Figure 6.**
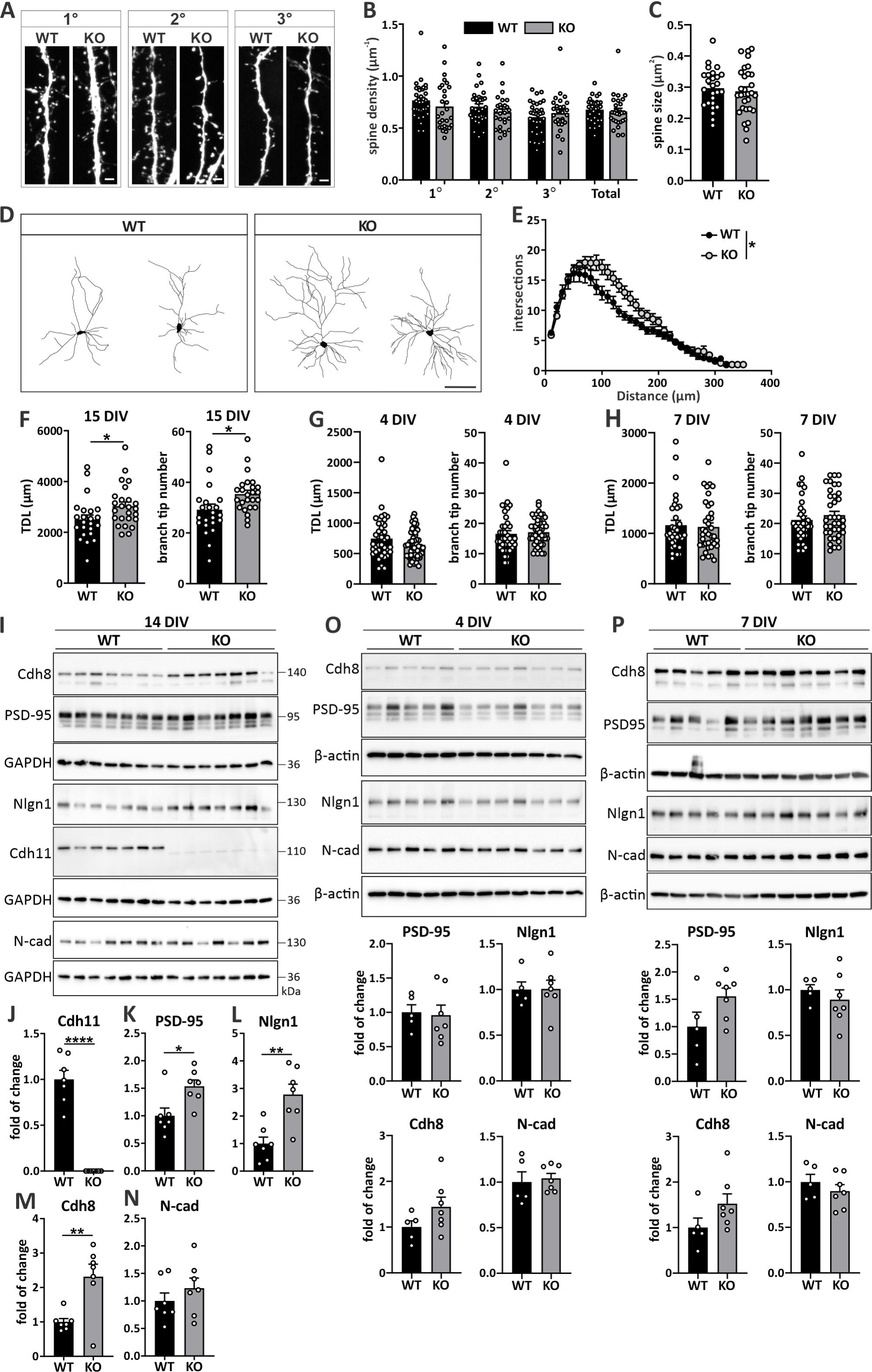
Increased dendrite complexity in *Cdh11^-/-^* neurons is accompanied by elevated expression levels of excitatory synaptic markers. (A) Confocal fluorochrome images of dendritic spines from primary, secondary and tertiary dendrites from 15 DIV *Cdh11* wild-type (WT) and knockout (KO) hippocampal neurons transfected with pLL3.7-GFP. Scale bar = 2 µm. **(B)** Quantification of the mean spine density (N = 33 WT and 28 KO neurons from 8 animals per genotype) and **(C)** mean spine size (26 WT neurons/7 animals and 30 KO neurons/8 animals) of 15 DIV *Cdh11* WT and KO hippocampal neurons. **(D)** Representative images of reconstructed dendritic trees from 15 DIV *Cdh11* WT and KO hippocampal neurons. Scale bar = 100 μm. **(E)** Sholl analysis of reconstructed neurons. Significant difference was determined by quantifying the area under the Sholl curve (AUC) between WT and KO neurons. \**p* < 0.05, unpaired two-tailed *t*-test. N = 23 WT and 25 KO neurons from 8 animals per genotype. Quantification of total dendrite length (TDL) and branch tip number of *Cdh11* WT and KO neurons at **(F)** 15 DIV, **(G)** 4 DIV and **(H)** 7 DIV. \**p* < 0.05, unpaired two-tailed *t*-test. 15 DIV: N = 23 WT and 25 KO neurons from 8 animals per genotype; 4 DIV: N = 38 WT neurons/4 animals and 52 KO neurons/3 animals; 7 DIV: N = 33 WT and 37 KO neurons from 4 animals per genotype. **(I)** Western blot analysis of the expression levels of cadherin-11, PSD-95, neuroligin-1, cadherin-8 and N-cadherin in 14 DIV *Cdh11* WT and KO hippocampal cultures. Cadherin-11 is not detectable in *Cdh11* KO cultures. Signals were normalized to GAPDH. Quantification of the expression of **(J)** cadherin-11, **(K)** PSD-95, **(L)** neuroligin-1, **(M)** cadherin-8 and **(N)** N-cadherin in 14 DIV *Cdh11* WT and KO hippocampal cultures. \**p <* 0.05, \*\**p <* 0.01, **** *p <* 0.0001, unpaired two-tailed *t*-test. N = 7 cultures per genotype. Expression profiles and quantification of PSD-95, neuroligin-1, cadherin-8 and N-cadherin in **(O)** 4 DIV and **(P)** 7 DIV *Cdh11* WT and KO hippocampal cultures. Signals were normalized to β-actin. N = 5 WT and 7 KO cultures.

### Expression levels of excitatory synaptic markers are increased in *Cdh11^-/-^* mice

Although synaptic density and spine size were not different in *Cdh11* knockout and wild-type mice, the changes in dendritic complexity suggested the possibility of an overall change of synaptic structures. Therefore, we examined protein levels of the excitatory synaptic marker PSD-95 in 14 DIV hippocampal cultures prepared from *Cdh11* wild-type and knockout mice. Western blot analysis confirmed no detectable expression of cadherin-11 in *Cdh11* knockout cultures (Figure 6I, J). However, PSD-95 levels were significantly increased in *Cdh11^-/-^* neurons (Figure 6I, K). We further examined protein levels of another synaptic protein, neuroligin-1, which is preferentially expressed at excitatory synapses (Song et al., 1999), and showed that neuroligin-1 was significantly increased in *Cdh11^-/-^* cultures compared to wild-type cultures (Figure 6I, L). Furthermore, we found similar expression levels of PSD-95 and neuroligin-1 in *Cdh11* knockout and wild-type neurons at both, 4 DIV (Figure 6O) and 7 DIV (Figure 6P), time points that did not show any changes in dendrite morphology (Figure 6G, H). This data suggests that the increase of dendrite complexity at a later developmental time point may likely result in an overall increase of postsynaptic sites in *Cdh11* knockout neurons without changing dendritic spine density or size.

Since we identified neuroligin-1 as a selective binding partner of cadherin-8, we reasoned that cadherin-8 levels may also be altered in *Cdh11^-/-^* neurons. Interestingly, cadherin-8 levels were not different at 4 and 7 DIV (Figure 6O, P), but were significantly increased in *Cdh11^-/-^* hippocampal neurons at 14 DIV (Figure 6I, M). For all time points analyzed, there was no difference in expression levels of the classical type I cadherin N-cadherin (Figure 6I, N-P). We also observed this altered protein expression profile of PSD-95, neuroligin-1 and cadherin-8 in brain lysates from *Cdh11* knockout and wild-type mice, suggesting that these changes are also detectable *in vivo* (Figure 7). However, in contrast to the cultured neurons, the changes in protein expression levels were readily apparent at an earlier time point *in vivo* (Figure 7A-D). Together, loss of cadherin-11 along with increased cadherin-8 expression in *Cdh11* knockout tissue recapitulates the expression profile observed in the NPCs from autistic human subjects (Figure 5) and further strengthens the confidence of using the *Cdh11* knockout mice to investigate potential mechanistic alterations occurring in autism.

**Figure 7.**
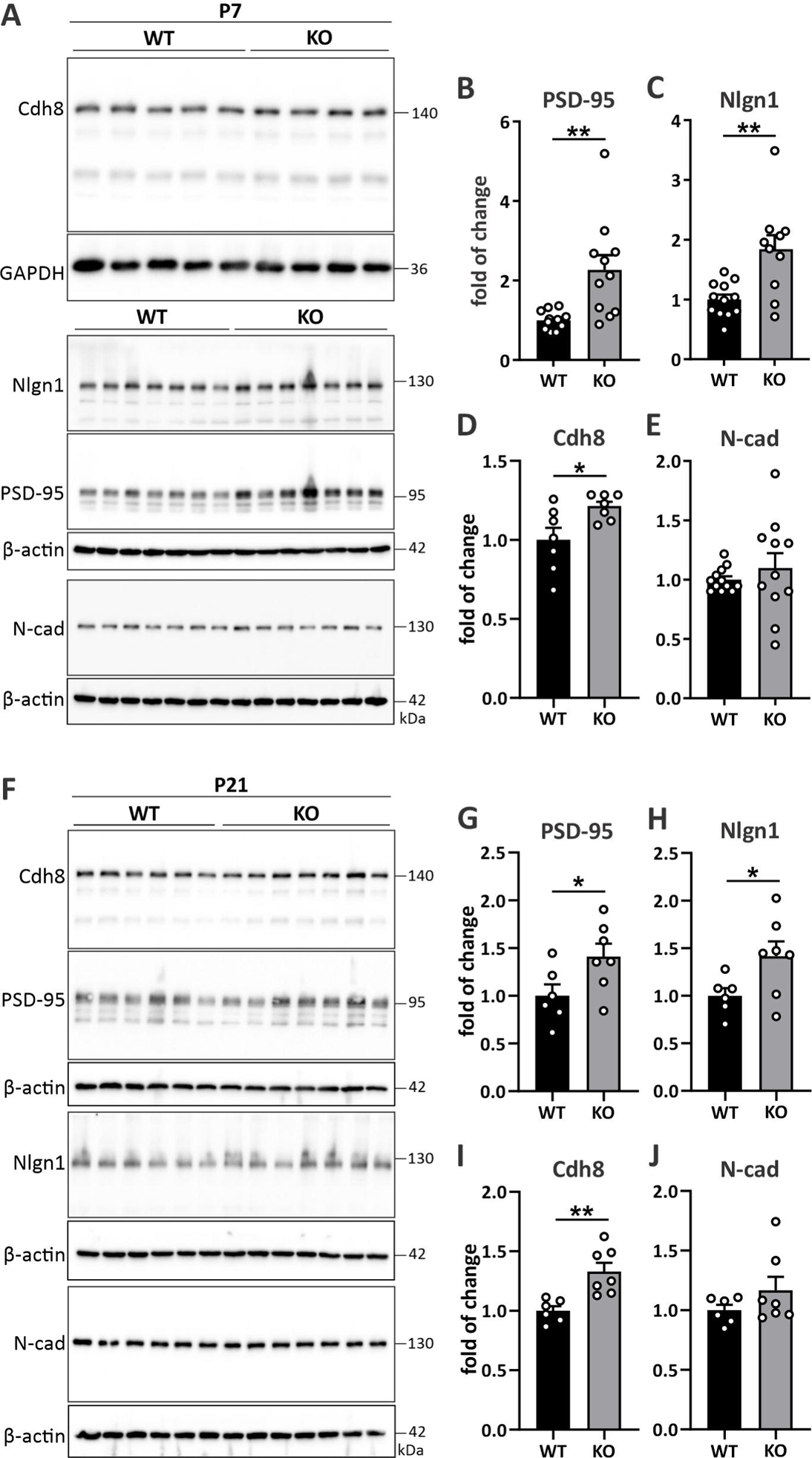
*Cdh11^-/-^* mice show elevated expression of PSD-95, neuroligin-1 and cadherin-8 *in vivo.* **(A)** Western blot analysis of PSD-95, neuroligin-1, cadherin-8 and N-cadherin expression in whole brains of P7 *Cdh11* WT and KO mice. Cadherin-8 signal was normalized to GAPDH and Nlgn1, PSD-95 and N-cadherin signals were normalized to β-actin. One representative immunoblot for Cdh8 with 5 WT and 4 KO samples is shown. Quantification of the expression of **(B)** PSD-95, **(C)** neuroligin-1, **(D)** cadherin-8 and **(E)** N-cadherin in P7 *Cdh11* WT and KO whole brains. \**p* < 0.05, \*\**p* < 0.01, unpaired two-tailed *t*-test. N = 7 animals per genotype (Cdh8), 12 WT and 10 KO (Nlgn1), 12 WT and 11 KO (PSD-95, N-cad). **(F)** Western blot analysis of PSD-95, neuroligin-1, cadherin-8 and N-cadherin expression in P21 forebrains of *Cdh11* WT and KO mice. Cadherin signals were normalized to β-actin. Quantification of the β expression of **(G)** PSD-95, **(H)** neuroligin-1, **(I)** cadherin-8 and **(J)** N-cadherin in P21 *Cdh11* WT and KO forebrains. \**p* < 0.05, \*\**p* < 0.01, unpaired two-tailed *t*-test. N = 6 WT and 7 KO animals.

### *Cdh11^-/-^* mice exhibit altered calcium activity and miniature excitatory postsynaptic currents

The increases of dendritic complexity and excitatory synaptic markers in *Cdh11^-/-^* neurons prompted us to investigate whether deletion of cadherin-11 results in changes of neuronal and synaptic activity. We first transduced hippocampal neurons prepared from *Cdh11* knockout and wild-type mice with the NeuroBurst Orange lentivirus (EssenBioscience Sartorius) and imaged network calcium activity using the IncuCyte S3 Live-Cell Analysis System for Neuroscience (EssenBioscience Sartorius) at 15 DIV (Figure 8A) for 24 hours. Interestingly, *Cdh11* knockout cultures showed significantly fewer active neurons compared to wild-type cultures (Figure 8D; WT: 1,756 ± 109.9, KO: 1,238 ± 159.1), although there was no difference in total cell number or infection rate between genotypes (Figure 8B, C; cell number - WT: 7,207 ± 1,431, KO: 7,454 ± 787.7, infection rate - WT: 0.5897 ± 0.0828, KO: 0.6452 ± 0.0719). In addition, the activity of *Cdh11^-/-^* neurons was significantly less correlated when compared to wild-type neurons (Figure 8E; WT: 1.405 ± 0.0681, KO: 1.111 ± 0.0657). *Cdh11^-/-^* neurons further exhibited significantly reduced mean burst strengths (Figure 8F; WT: 0.6473 ± 0.0507, KO: 0.4163 ± 0.0324), whereas mean burst rate was similar in *Cdh11* knockout and wild-type cultures (Figure 8G; WT: 3.643 ± 0.9519 min^-1^, KO: 4.983 ± 1.424 min^-1^). Together, these data show that *Cdh11* knockout neurons exhibit a significant reduction in neuronal activity and synchrony compared to wild-type neurons, which suggests that synaptic input is reduced or altered.

**Figure 8.**
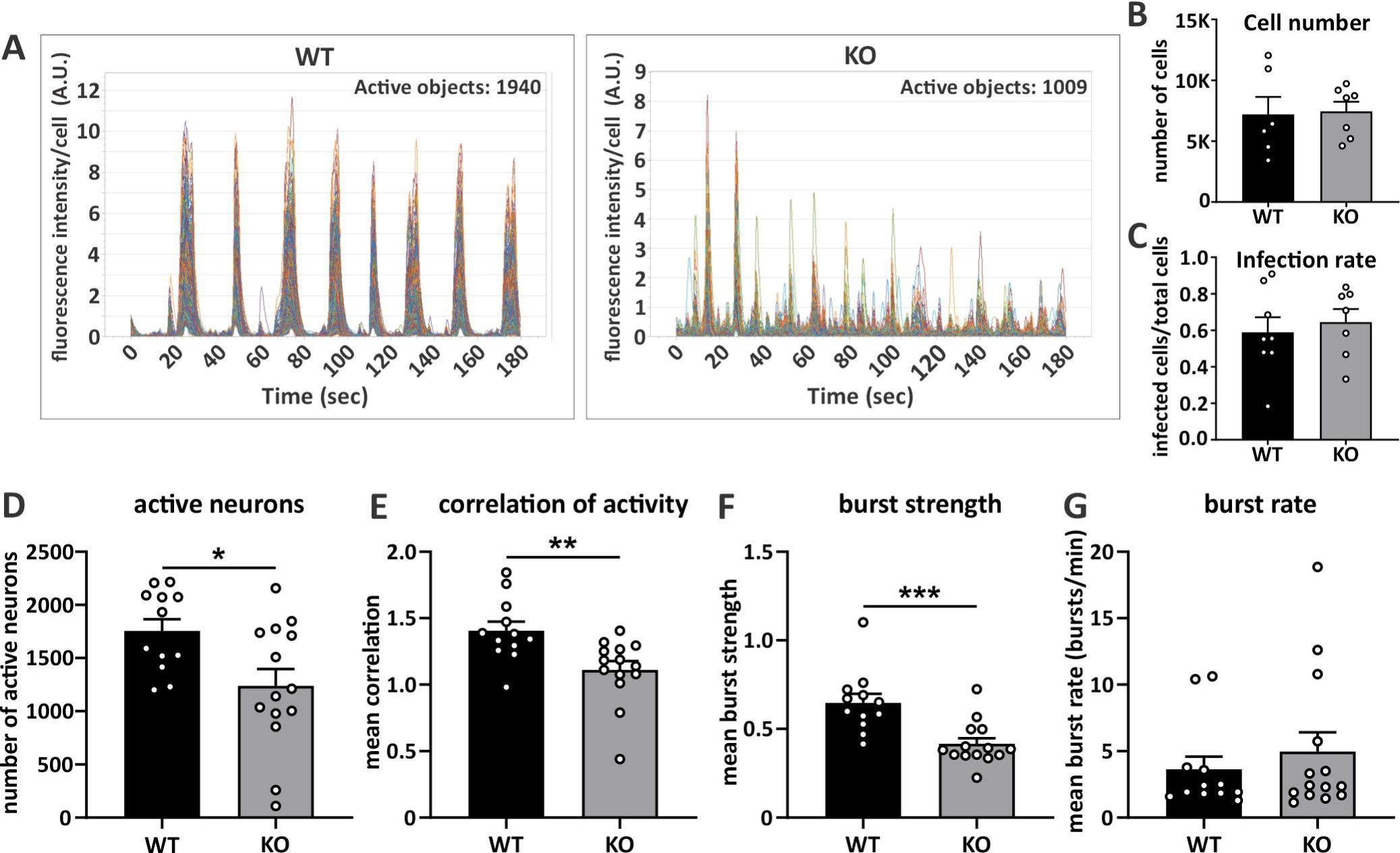
*Cdh11^-/-^* neurons show altered calcium activity. **(A)** Example calcium current traces from *Cdh11* WT and KO hippocampal cultures at 15 DIV recorded for 3 minutes using NeuroBurst Orange lentivirus calcium indicator. Averaged fluorescence intensity per cell of 1940 active neurons (WT) and 1009 active neurons (KO) was plotted against time (sec). **(B)** Total cell count of *Cdh11* WT and KO cultures using DAPI counterstain at 16 DIV. N = 6 WT and 7 KO cultures. **(C)** Infection rate of *Cdh11* WT and KO neurons at 16 DIV infected by the IncuCyte Neurolight Orange lentivirus. The number of infected cells was normalized to the total number of neurons counterstained with DAPI. N = 8 WT and 7 KO cultures. Measurement of **(D)** number of active neurons, **(E)** correlation of activity, **(F)** mean burst strength and (**G)** mean burst rate of *Cdh11* WT and KO hippocampal neurons infected with the NeuroBurst Orange lentivirus calcium indicator recorded at 15 DIV and 16 DIV. \**p* < 0.05, \*\**p* < 0.01, \*\*\**p* < 0.001, unpaired two-tailed *t*-test. N = 6 WT and 7 KO cultures from two different time points.

Because of these activity deficits, we hypothesized that excitatory synaptic input is diminished by cadherin-11 deletion. We therefore investigated hippocampal synaptic function in animals at two developmental time points. We used *ex vivo* hippocampal slices to examine the effects of *Cdh11* knockout on AMPAR-mediated miniature excitatory postsynaptic currents (mEPSCs) in CA1 pyramidal cells from wild-type and knockout mice aged P7-P9 (Figure 9A, B) and P21-P24 (Figure 9H, I). We did not observe any differences in miniature EPSC frequency and amplitude in CA1 pyramidal cells from *Cdh11* knockout and wild-type mice aged P7-P9 (Figure 9C, D; frequency - WT: 0.1669 ± 0.0221 Hz, KO: 0.1891 ± 0.0308 Hz; amplitude - WT: 12.73 ± 0.9390 pA, KO: 13.07 ± 0.5904 pA). Also, no differences in mEPSC kinetics were observed between P7-P9 *Cdh11* knockout and wild-type mice (Figure 9E-G; charge - WT: 87.32 ± 7.684 fC, KO: 83.74 ± 4.505 fC; rise time - WT: 1.246 ± 0.0.0938 ms, KO: 1.180 ± 0.0805 ms; decay time - WT: 4.643 ± 0.3093 ms, KO: 4.533 ± 0.2207 ms). However, mEPSC frequency was significantly lower in CA1 pyramidal cells from *Cdh11* knockout mice compared to wild-type littermates at the later time point in development, i.e. P21-P24 (Figure 9J; WT: 1.16 ± 0.1254 Hz, KO: 0.5026 ± 0.0445 Hz). Conversely, mEPSC amplitude was higher in *Cdh11* knockout neurons (Figure 9K; WT: 11.28 ± 0.4684 pA, KO: 14.12 ± 1.078 pA). Analysis of the kinetics of CA1 mEPSCs found that there was no difference in the charge (Figure 9L; WT: 76.92 ± 4.035 fC, KO: 91.56 ± 6.325 fC), rise time (Figure 9M; WT: 1.355 ± 0.0444 ms, KO: 1.355 ± 0.0644 ms) or decay time constant between P21-P24 *Cdh11* knockout and wild-type mice (Figure 9N; WT: 4.940 ± 0.1944 ms, KO: 5.091 ± 0.2281 ms). The sharp reduction of mEPSC frequency in P21-P24 *Cdh11^-/-^* CA1 neurons most likely reflects a reduction in presynaptic function or the number of functional presynaptic release sites. The concurrent increase in mEPSC amplitude is consistent with the overall increased expression of the postsynaptic markers PSD-95 and neuroligin-1. Together, these results suggest that loss of a trans-synaptic protein, such as cadherin-11, disrupts typical synaptic organization and function, and therefore neuronal activity.

**Figure 9.**
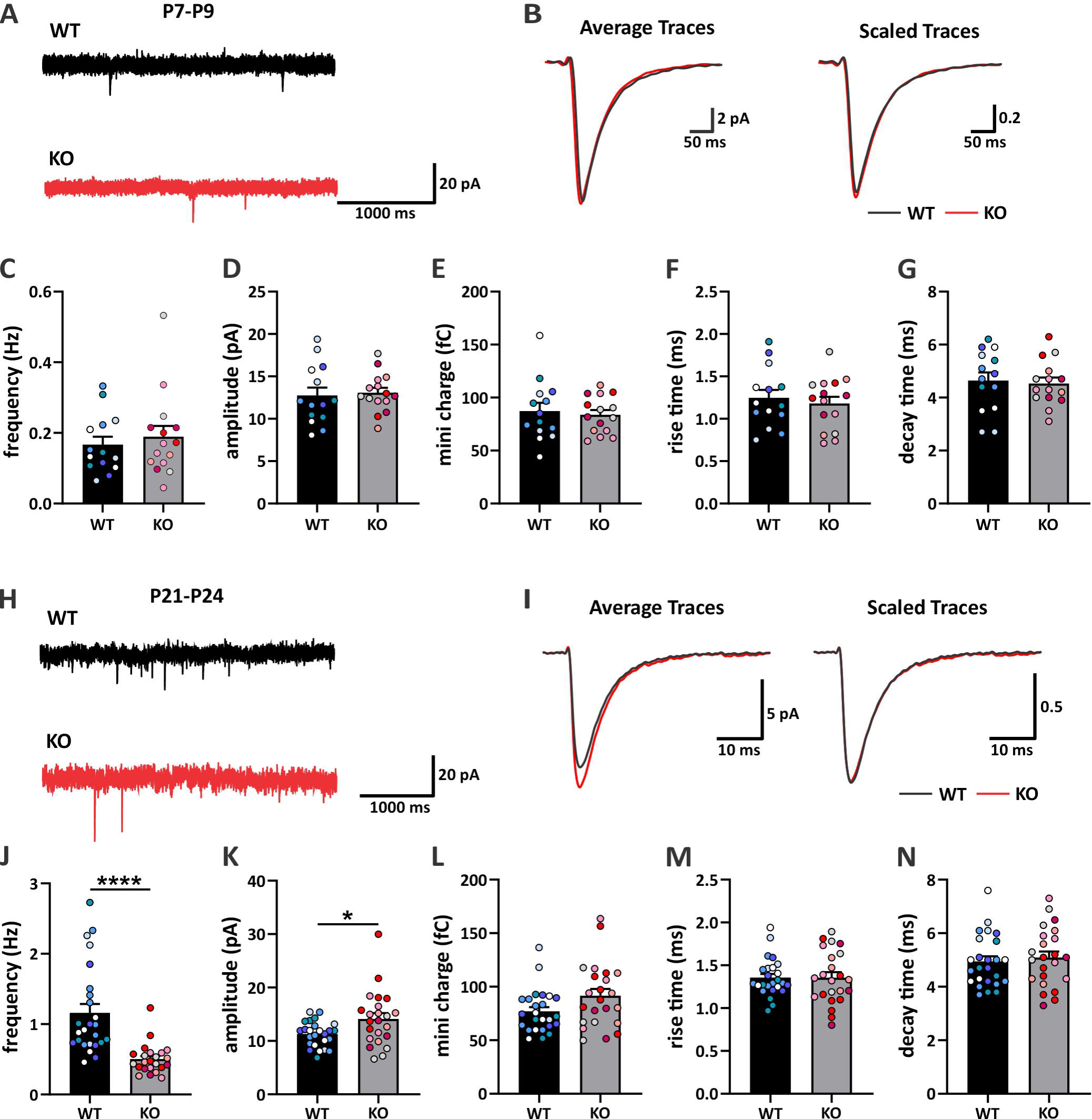
*Cdh11^-/-^* mice exhibit changes in miniature excitatory postsynaptic currents. **(A)** Example current traces recorded from whole-cell voltage-clamp of hippocampal CA1 pyramidal neurons from P7-P9 *Cdh11* WT and KO mice. **(B)** Examples of average (left) and scaled (right) mEPSC traces from P7-P9 *Cdh11* WT and KO cells. Measurement of **(C)**frequency, **(D)** amplitude, **(E)** charge, **(F)** rise time and **(G)** decay time constant of recorded mEPSCs. N = 14 WT and 15 KO neurons from 5 animals per genotype. **(H)** Example current traces recorded from whole-cell voltage-clamp of hippocampal CA1 pyramidal neurons from P21-P24 mice. **(I)** Examples of average (left) and scaled (right) mEPSC traces from P21-P24 *Cdh11* WT and KO cells. Measurement of **(J)** frequency, **(K)** amplitude, **(L)** charge, **(M)** rise time and **(N)** decay time constant of recorded mEPSCs. \**p* < 0.05, \*\*\*\**p* < 0.0001, unpaired two-tailed *t*-test. N = 25 WT and 23 KO neurons from 5 animals per genotype. Color-codes indicate individual data points from one animal.

Our findings from neuronal cultures and animals together indicate that loss of cadherin-11 results in altered dendritic morphology, changes in expression of excitatory synaptic markers, and disturbed synaptic transmission. These phenotypes may likely underlie neuronal alterations observed in autism.

## Discussion

In this study, we first evaluated the potential roles of the autism candidate risk genes CDH8 and CDH11 by systematically investigating their expression and localization in mouse tissue. Comprehensive analysis revealed that both of these type II classical cadherins are expressed in the developing mouse brain at the necessary time and places to regulate crucial steps in neural circuit formation and to impact normal brain development. Further, expression analysis of CDH8 and CDH11 in NPCs and mature organoids generated from autistic individuals demonstrated that levels of the two cadherins are altered in human conditions. The analysis of the *Cdh11^-/-^* mouse neurons further uncovered cellular and circuitry phenotypes that may be observed in autism. Our study not only describes a detailed characterization of the autism-risk genes CDH8 and CDH11, but also provides insights into novel functions of cadherin-11 in neural circuit development that may potentially be implicated in the etiology of autism.

Cadherin-8 and cadherin-11 belong to the same classical type II cadherin subfamily. In addition to a similar structure, several studies have found similar expression profiles and functions of these two cadherins in the brain. Cadherin-8 is important for synaptic targeting in the hippocampus (Bekirov et al., 2008), striatum (Friedman et al., 2015b), and visual system (Osterhout et al., 2011; Duan et al., 2014). Cadherin-11 strongly expresses in the hippocampus and functions at glutamatergic synapses (Manabe et al., 2000; Paradis et al., 2007; Bartelt-Kirbach et al., 2010). In agreement with these studies, our spatial expression analysis shows that cadherin-8 and cadherin-11 are enriched in similar brain regions including cortex, hippocampus, and thalamus/striatum. The temporal expression analysis shows that both cadherins exhibit peak expression in postnatal brains at the time window that coincides with dendrite development and synaptogenesis, processes that have been adversely implicated in the developing autism brain (Betancur et al., 2009; Bourgeron, 2009; Hussman et al., 2011). In particular, cadherin-8 exhibited a dramatic but transient increase from embryonic stage to the first postnatal week. Protein levels quickly declined later in development and remained low in adulthood. This surge in expression indicates a putative role for cadherin-8 in regulating dendrite and synapse development, which is further supported by cadherin-8 localization to synaptic puncta. Although a direct interaction can be demonstrated between cadherin-8 and cadherin-11, we found neuroligin-1 as a selective interaction partner of cadherin-8, but not of cadherin-11, suggesting the presence of partially divergent signaling pathways and functions mediated by these two closely related cadherins. Therefore, the increase of cadherin-8 levels in the absence of cadherin-11, potentially due to compensation, may result in changes of the selective interaction between cadherin-8 and neuroligin-1 and contribute to the altered cellular and physiological phenotypes. Future studies should address a direct effect of cadherin-8, neuroligin-1, PSD95, or other proteins that may be up- or downregulated in the *Cdh11* knockout mice to clarify their roles in the observed neuronal phenotypes.

The enrichment of cadherin-11 in synaptic plasma membrane fractions suggests localization on both sides of the synapse, where cadherin-11 may span the synaptic cleft to act as a synaptic cell adhesion molecule. Deletion of cadherin-11 could impact the interaction between pre- and postsynaptic sites, which may result in disturbed synaptic function. In line with this hypothesis is the dramatic reduction in mEPSC frequency in P21-P24 *Cdh11* knockout neurons, which may indicate deficits in presynaptic site number and/or function, or neurotransmitter vesicle release. The concurrent increase of mEPSC amplitude is compatible with the increased PSD-95 and neuroligin-1 protein levels and could represent a compensatory mechanism of the *Cdh11* knockout neurons to reach homeostatic levels of synaptic input, or could be a direct effect of loss of cadherin-11 disrupting synaptic organization, a possibility that is consistent with the absence of an increase in dendritic spine size. The reduction in mEPSC frequency is also compatible with the observed reduction in calcium-reporter activity, as indicated by the decreased correlation of activity and lower proportion of active cells, as well as reduced mean burst strengths, reflecting an overall decrease in somatic calcium influx that could arise from diminished synaptic input. The increase in dendritic arborization may reflect structural homeostasis, where *Cdh11^-/-^* neurons adjust the dendritic arbors and search for more synaptic input in response to reduced neuronal activity (Tripodi et al., 2008). Future studies are required to determine if the altered morphology *in vitro* is also observed *in vivo*, where the contribution of brain area and cell type must be additionally considered.

An alternative explanation for reduced mEPSC frequency is an increased number of silent synapses. In addition to the synaptic plasma membrane, cadherin-11 was also enriched in postsynaptic density fractions, where it may be required for anchoring or organizing receptors. Thus, depletion of cadherin-11 may impact postsynaptic AMPA-receptor expression, and consequently affect the proportion of non-functional “deaf” synapses (Voronin & Cherubini, 2004). Likewise, loss of cadherin-11, as a presynaptic adhesion molecule, could disrupt pre-synaptic organization or trans-synaptic connections, leading to “mute” synapses (Crawford and Mennerick, 2012). Further investigations are required to help elucidate a potential mechanism underlying the altered synaptic activity in *Cdh11*knockout brains, including more detailed analysis of the presynaptic terminals, such as number, location and co-localization with PSD-95 puncta, as well as analysis of the postsynaptic site, such as number and distribution of AMPA-receptors between synapses.

Members of the cadherin superfamily have emerged as candidate risk genes for autism in multiple independent association studies (Marshall et al., 2008; Morrow et al., 2008; Depienne et al., 2009; Wang et al., 2009; Willemsen et al., 2010; Chapman et al., 2011; Hussman et al., 2011; Pagnamenta et al., 2011; Sanders et al., 2011; Camacho et al., 2012; Neale et al., 2012; O’Roak et al., 2012; Girirajan et al., 2013; van Harssel et al., 2013; Crepel et al., 2014; Cukier et al., 2014; Kenny et al., 2014). Despite these findings, no autism case reported to date has resulted from monogenic alterations in cadherins. In fact, none of the individuals with autism tested in this study harbor damaging mutations in the *CDH8* or *CDH11* genes (DeRosa et al., 2018). Yet, these individuals consistently show altered expression patterns in CDH8 and CDH11 throughout neural development from neural precursor cells to more mature organoids. Despite the genetic and clinical heterogeneity seen in ASD, which is reflected by some inter-individual variability shown in the expression of these cadherins in iPSC-derived NPCs, the significant differences in CDH8 and CDH11 expression suggest a potential vulnerable mechanism that involves these two cadherins. In addition to specific autism-associated mutations in cadherins, these molecules may be part of a broader pathway involving other genes mutated in autism. Indeed, some of the autism lines examined in this study also display misregulation of genes involved in cell-cell signaling and actin cytoskeleton signaling, both processes that involve cadherins (DeRosa et al., 2018). The overall decrease in synaptic activity that we observed in the *Cdh11* knockout neurons is consistent with the phenotypic characterization of the iPSC-derived cortical neurons from autistic individuals performed by DeRosa et al., as they reported decreased spontaneous spiking activity, as well as decreased number of calcium transients in these neurons. The autism-specific iPSC-derived cortical neurons also exhibited decreased migration of neuronal processes (DeRosa et al., 2018). Interestingly, elevated expression of CDH11 has been observed in glioblastoma and has been associated with increased cell migration (Schulte et al., 2013). Thus, the reduced migration ability of the autism-specific neurons may stem from the decreased CDH11 expression that we observed in these autism lines. These findings further emphasize the significance of our effort to investigate the involvement of CDH11 in autism.

The variants in the CDH8 and CDH11 genes identified in individuals with autism are predicted to cause loss of function (Pagnamenta et al., 2011, Crepel et al., 2014). We found downregulation of CDH11 in autism-specific NPCs, which could mimic a loss-of-function phenotype. In contrast, NPCs from autistic individuals showed upregulation of CDH8, which may be at odds with the autism-associated loss-of-function mutations in the CDH8 gene. However, based on the known functions of cadherins, both lower and higher expression levels could directly affect neuronal connectivity by altering cell-cell adhesion, downstream signaling, and synaptic function. The autism cases investigated in this study did not carry any genetic alterations in CDH8 and CDH11 genes, and yet, they exhibited changes in protein levels of these two cadherins, further strengthening the hypothesis that cadherin signaling may represent an important pathway affected in autism. Upstream factors affecting the expression of cadherin-8 and cadherin-11 still require further investigation. Future studies should also aim to identify mechanisms that connect different cadherins to other synaptic molecules. An example of a potential connection is described here between cadherin-8 and neuroligin-1, another autism risk candidate molecule (Nakanishi et al., 2017). In conclusion, our study shows that cadherin-11 plays an important role in regulating dendritic morphology and synaptic function of excitatory neurons, and that altered expression of cadherin-11 may contribute in part to phenotypes observed in autism.

### Data availability statement

The dataset supporting the conclusions of this study are available upon reasonable request by the corresponding authors.

## Acknowledgements

We want to thank Dr. Celine Plachez at the Hussman Institute for Autism for the technical support. We also want to thank Drs. Louis DeTolla and Turhan Coksaygan at the University of Maryland School of Medicine for providing veterinary services and consultation. We thank Dr. Olga Latinovic from the imaging core at the Institute of Human Virology at the University of Maryland School of Medicine for her help and assistance with the dSTORM imaging. Finally, we thank Drs. John Hussman, Anthony Koleske, Thomas Blanpied, and Deanna Benson for the critical evaluation of this manuscript.

## Author contributions

JF, RN, MB, MD, MK, XY, DD, MN, SH, GB, YL Designed research; JF, RN, MB, MD, JN, MK Performed research; JF, RN, MB, MD, MK, YL Analyzed data; JF, RN, MB, MD, DD, MN, SH, GB, YL Wrote paper

---

**Figure 1-1. Specificity of antibodies. (A)** Specificity of antibodies was determined by Western blot (WB). WB of untransfected N2a cells, N2a cells transfected (TF) with myc-flag-tagged *Cdh8* or flag-tagged *Cdh11* and P14 mouse brain tissues were probed with mouse anti-Cdh8, goat anti-Cdh8 or mouse anti-Cdh11 antibodies. Each antibody recognizes prominent bands of predicted molecular weight (Cdh8: 140 kDa (precursor), 90 kDa (mature); Cdh11: 110 kDa), each band representing either the overexpressed protein in N2a cells or the endogenous protein in brain tissues. Note that Cdh8 and Cdh11 are not endogenously expressed in N2a cells. **(B)** Cross-reactivity of antibodies. WB of N2a cells transfected with myc-flag-tagged *Cdh8* or flag-tagged *Cdh11* were probed with the same antibodies as in (A). Each antibody specifically recognizes the predicted cadherin and is not cross-reacting with the other cadherin examined. GAPDH served as loading control. **(C, D)** Specificity of antibodies tested by immunofluorescence. **(C)** myc-flag-tagged *Cdh8-* or **(D)** HA-tagged *Nlgn-1*-transfected N2a cells were fixed and stained with the mouse anti-Cdh8 or mouse anti-Nlgn-1 antibody. These antibodies specifically recognize the overexpressed cadherin-8 (cyan) and its flag-tag (magenta) and the overexpressed neuroligin-1 (cyan) and its HA-tag (magenta), respectively, with characteristic localization to cell membranes and cell-cell contacts. No signals were detectable in untransfected N2a cells that were incubated with primary and secondary antibodies. DAPI signal is depicted in blue. Scale bar = 10 μm (C), 20 μm (D).

## References

American Psychiatric Association (2013). Diagnostic and Statistical Manual of Mental Disorders, 5th Edn. Arlington, VA: American Psychiatric Publishing

Aiga M, Levinson JN, Bamji SX (2011) N-cadherin and neuroligins cooperate to regulate synapse formation in hippocampal cultures. J Biol Chem 286:851–858.

Arikkath J, Reichardt LF (2008) Cadherins and catenins at synapses: roles in synaptogenesis and synaptic plasticity. Trends Neurosci 31:487–494.

Bartelt-Kirbach B, Langer-Fischer K, Golenhofen N (2010) Different regulation of N-cadherin and cadherin-11 in rat hippocampus. Cell Commun Adhes 17:75–82.

Basu R, Taylor MR, Williams ME (2015) The classic cadherins in synaptic specificity. Cell Adh Migr 9:193–201.

Bekirov IH, Nagy V, Svoronos A, Huntley GW, Benson DL (2008) Cadherin-8 and N-cadherin differentially regulate pre- and postsynaptic development of the hippocampal mossy fiber pathway. Hippocampus 18:349–363.

Bermejo MK, Milenkovic M, Salahpour A, Ramsey AJ (2014) Preparation of synaptic plasma membrane and postsynaptic density proteins using a discontinuous sucrose gradient. J Vis Exp:e51896.

Betancur C, Sakurai T, Buxbaum JD (2009) The emerging role of synaptic cell-adhesion pathways in the pathogenesis of autism spectrum disorders. Trends Neurosci 32:402–412.

Bourgeron T (2009) A synaptic trek to autism. Curr Opin Neurobiol 19:231–234.

Bourgeron T (2015) From the genetic architecture to synaptic plasticity in autism spectrum disorder. Nat Rev Neurosci 16:551–563.

Brasch J, Katsamba PS, Harrison OJ, Ahlsén G, Troyanovsky RB, Indra I, Kaczynska A, Kaeser B, Troyanovsky S, Honig B, Shapiro L (2018) Homophilic and Heterophilic Interactions of Type II Cadherins Identify Specificity Groups Underlying Cell-Adhesive Behavior. Cell Rep 23:1840–1852.

Camacho A, Simón R, Sanz R, Viñuela A, Martínez-Salio A, Mateos F (2012) Cognitive and behavioral profile in females with epilepsy with PDCH19 mutation: two novel mutations and review of the literature. Epilepsy Behav 24:134–137.

Chapman NH, Estes A, Munson J, Bernier R, Webb SJ, Rothstein JH, Minshew NJ, Dawson G, Schellenberg GD, Wijsman EM (2011) Genome-scan for IQ discrepancy in autism: evidence for loci on chromosomes 10 and 16. Hum Genet 129:59–70.

Chen J, Yu S, Fu Y, Li X (2014) Synaptic proteins and receptors defects in autism spectrum disorders. Front Cell Neurosci 8:276.

Chih B, Gollan L, Scheiffele P (2006) Alternative splicing controls selective trans-synaptic interactions of the neuroligin-neurexin complex. Neuron 51:171–178.

Crawford DC, Mennerick S (2012) Presynaptically silent synapses: dormancy and awakening of presynaptic vesicle release. Neuroscientist, 18:216–223.

Crepel A, De Wolf V, Brison N, Ceulemans B, Walleghem D, Peuteman G, Lambrechts D, Steyaert J, Noens I, Devriendt K, Peeters H (2014) Association of CDH11 with non-syndromic ASD. Am J Med Genet B Neuropsychiatr Genet 165B:391–398.

Cukier HN, Dueker ND, Slifer SH, Lee JM, Whitehead PL, Lalanne E, Leyva N, Konidari I, Gentry RC, Hulme WF, Booven DV, Mayo V, Hofmann NK, Schmidt MA, Martin ER, Haines JL, Cuccaro ML, Gilbert JR, Pericak-Vance MA (2014) Exome sequencing of extended families with autism reveals genes shared across neurodevelopmental and neuropsychiatric disorders. Mol Autism 5:1.

Depienne C et al. (2009) Sporadic infantile epileptic encephalopathy caused by mutations in PCDH19 resembles Dravet syndrome but mainly affects females. PLoS Genet 5:e1000381.

DeRosa BA, Van Baaren JM, Dubey GK, Lee JM, Cuccaro ML, Vance JM, Pericak-Vance MA, Dykxhoorn DM (2012) Derivation of autism spectrum disorder-specific induced pluripotent stem cells from peripheral blood mononuclear cells. Neurosci Lett 516:9–14.

DeRosa BA, El Hokayem J, Artimovich E, Garcia-Serje C, Phillips AW, Van Booven D, Nestor JE, Wang L, Cuccaro ML, Vance JM, Pericak-Vance MA, Cukier HN, Nestor MW, Dykxhoorn DM (2018) Convergent Pathways in Idiopathic Autism Revealed by Time Course Transcriptomic Analysis of Patient-Derived Neurons. Sci Rep 8:8423.

Duan X, Krishnaswamy A, De la Huerta I, Sanes JR (2014) Type II cadherins guide assembly of a direction-selective retinal circuit. Cell 158:793–807.

Durens M, Nestor J, Williams M, Herold K, Niescier RF, Lunden JW, Phillips AW, Lin YC, Dykxhoorn DM, Nestor MW (2020) High-throughput screening of human induced pluripotent stem cell-derived brain organoids. J Neurosci Methods 335:108627.

Ferreira TA, Blackman AV, Oyrer J, Jayabal S, Chung AJ, Watt AJ, Sjöström PJ, van Meyel DJ (2014) Neuronal morphometry directly from bitmap images. Nat Methods 11:982–984.

Forrest MP, Parnell E, Penzes P (2018) Dendritic structural plasticity and neuropsychiatric disease. Nat Rev Neurosci 19:215–234.

Friedman LG, Benson DL, Huntley GW (2015a) Cadherin-based transsynaptic networks in establishing and modifying neural connectivity. Curr Top Dev Biol 112:415–465.

Friedman LG, Riemslagh FW, Sullivan JM, Mesias R, Williams FM, Huntley GW, Benson DL (2015b) Cadherin-8 expression, synaptic localization, and molecular control of neuronal form in prefrontal corticostriatal circuits. J Comp Neurol 523:75–92.

Girirajan S, Dennis MY, Baker C, Malig M, Coe BP, Campbell CD, Mark K, Vu TH, Alkan C, Cheng Z, Biesecker LG, Bernier R, Eichler EE (2013) Refinement and discovery of new hotspots of copy-number variation associated with autism spectrum disorder. Am J Hum Genet 92:221–237.

Gu Y, Huganir RL (2016) Identification of the SNARE complex mediating the exocytosis of NMDA receptors. Proc Natl Acad Sci U S A 113:12280–12285.

Hirano S, Takeichi M (2012) Cadherins in brain morphogenesis and wiring. Physiol Rev 92:597–634.

Horikawa K, Radice G, Takeichi M, Chisaka O (1999) Adhesive subdivisions intrinsic to the epithelial somites. Dev Biol 215:182–189.

Huguet G, Ey E, Bourgeron T (2013) The genetic landscapes of autism spectrum disorders. Annu Rev Genomics Hum Genet 14:191–213.

Hulpiau P, van Roy F (2009) Molecular evolution of the cadherin superfamily. Int J Biochem Cell Biol 41:349–369.

Hussman JP, Chung RH, Griswold AJ, Jaworski JM, Salyakina D, Ma D, Konidari I, Whitehead PL, Vance JM, Martin ER, Cuccaro ML, Gilbert JR, Haines JL, Pericak-Vance MA (2011) A noise-reduction GWAS analysis implicates altered regulation of neurite outgrowth and guidance in autism. Mol Autism 2:1.

Hutsler JJ, Zhang H (2010) Increased dendritic spine densities on cortical projection neurons in autism spectrum disorders. Brain Res 1309:83–94.

Inoue T, Tanaka T, Suzuki SC, Takeichi M (1998) Cadherin-6 in the developing mouse brain: expression along restricted connection systems and synaptic localization suggest a potential role in neuronal circuitry. Dev Dyn 211:338–351.

Jeste SS, Geschwind DH (2014) Disentangling the heterogeneity of autism spectrum disorder through genetic findings. Nat Rev Neurol 10:74–81.

Joensuu M, Lanoue V, Hotulainen P (2017) Dendritic spine actin cytoskeleton in autism spectrum disorder. Prog Neuropsychopharmacol Biol Psychiatry.

Kenny EM, Cormican P, Furlong S, Heron E, Kenny G, Fahey C, Kelleher E, Ennis S, Tropea D, Anney R, Corvin AP, Donohoe G, Gallagher L, Gill M, Morris DW (2014) Excess of rare novel loss-of-function variants in synaptic genes in schizophrenia and autism spectrum disorders. Mol Psychiatry 19:872–879.

Korematsu K, Redies C (1997) Expression of cadherin-8 mRNA in the developing mouse central nervous system. J Comp Neurol 387:291–306.

Lin YC, Frei JA, Kilander MB, Shen W, Blatt GJ (2016) A Subset of Autism-Associated Genes Regulate the Structural Stability of Neurons. Front Cell Neurosci 10:263.

Lord C, Rutter M, Le Couteur A (1994) Autism Diagnostic Interview-Revised: a revised version of a diagnostic interview for caregivers of individuals with possible pervasive developmental disorders. J Autism Dev Disord 24:659–685.

Maenner MJ et al. (2020) Prevalence of Autism Spectrum Disorder Among Children Aged 8 Years - Autism and Developmental Disabilities Monitoring Network, 11 Sites, United States, 2016. MMWR Surveill Summ 69:1–12.

Manabe T, Togashi H, Uchida N, Suzuki SC, Hayakawa Y, Yamamoto M, Yoda H, Miyakawa T, Takeichi M, Chisaka O (2000) Loss of cadherin-11 adhesion receptor enhances plastic changes in hippocampal synapses and modifies behavioral responses. Mol Cell Neurosci 15:534–546.

Marshall CR et al. (2008) Structural variation of chromosomes in autism spectrum disorder. Am J Hum Genet 82:477–488.

Meijering E, Jacob M, Sarria JC, Steiner P, Hirling H, Unser M (2004) Design and validation of a tool for neurite tracing and analysis in fluorescence microscopy images. Cytometry A 58:167–176.

Morrow EM et al. (2008) Identifying autism loci and genes by tracing recent shared ancestry. Science 321:218–223.

Nahidiazar L, Agronskaia AV, Broertjes J, van den Broek B, Jalink K (2016) Optimizing Imaging Conditions for Demanding Multi-Color Super Resolution Localization Microscopy. PLoS One 11:e0158884.

Nakanishi M, Nomura J, Ji X, Tamada K, Arai T, Takahashi E, Bu an M, Takumi T (2017) Correction: Functional significance of rare neuroligin 1 variants found in autism. PLoS Genet 13:e1007035.

Neale BM et al. (2012) Patterns and rates of exonic de novo mutations in autism spectrum disorders. Nature 485:242–245.

Nestor MW, Jacob S, Sun B, Prè D, Sproul AA, Hong SI, Woodard C, Zimmer M, Chinchalongporn V, Arancio O, Noggle SA (2015) Characterization of a subpopulation of developing cortical interneurons from human iPSCs within serum-free embryoid bodies. Am J Physiol Cell Physiol 308:C209–219.

Noel J, Ralph GS, Pickard L, Williams J, Molnar E, Uney JB, Collingridge GL, Henley JM (1999) Surface expression of AMPA receptors in hippocampal neurons is regulated by an NSF-dependent mechanism. Neuron 23:365–376.

O’Roak BJ et al. (2012) Sporadic autism exomes reveal a highly interconnected protein network of de novo mutations. Nature 485:246–250.

Osterhout JA, Josten N, Yamada J, Pan F, Wu SW, Nguyen PL, Panagiotakos G, Inoue YU, Egusa SF, Volgyi B, Inoue T, Bloomfield SA, Barres BA, Berson DM, Feldheim DA, Huberman AD (2011) Cadherin-6 mediates axon-target matching in a non-image-forming visual circuit. Neuron 71:632–639.

Ovesný M, K ížek P, Borkovec J, Svindrych Z, Hagen GM (2014) ThunderSTORM: a comprehensive ImageJ plug-in for PALM and STORM data analysis and super-resolution imaging. Bioinformatics 30:2389–2390.

Pagnamenta AT, Khan H, Walker S, Gerrelli D, Wing K, Bonaglia MC, Giorda R, Berney T, Mani E, Molteni M, Pinto D, Le Couteur A, Hallmayer J, Sutcliffe JS, Szatmari P, Paterson AD, Scherer SW, Vieland VJ, Monaco AP (2011) Rare familial 16q21 microdeletions under a linkage peak implicate cadherin 8 (CDH8) in susceptibility to autism and learning disability. J Med Genet 48:48–54.

Paradis S, Harrar DB, Lin Y, Koon AC, Hauser JL, Griffith EC, Zhu L, Brass LF, Chen C, Greenberg ME (2007) An RNAi-based approach identifies molecules required for glutamatergic and GABAergic synapse development. Neuron 53:217–232.

Phillips AW, Nestor JE, Nestor MW (2017) Developing HiPSC Derived Serum Free Embryoid Bodies for the Interrogation of 3-D Stem Cell Cultures Using Physiologically Relevant Assays. J Vis Exp.

Redies C, Hertel N, Hübner CA (2012) Cadherins and neuropsychiatric disorders. Brain Res 1470:130–144.

Rubinson DA, Dillon CP, Kwiatkowski AV, Sievers C, Yang L, Kopinja J, Rooney DL, Zhang M, Ihrig MM, McManus MT, Gertler FB, Scott ML, Van Parijs L (2003) A lentivirus-based system to functionally silence genes in primary mammalian cells, stem cells and transgenic mice by RNA interference. Nat Genet 33:401–406.

Sanders SJ et al. (2011) Multiple recurrent de novo CNVs, including duplications of the 7q11.23 Williams syndrome region, are strongly associated with autism. Neuron 70:863–885.

Schulte JD, Srikanth M, Das S, Zhang J, Lathia JD, Yin L, Rich JN, Olson EC, Kessler JA, Chenn A (2013) Cadherin-11 regulates motility in normal cortical neural precursors and glioblastoma. PLoS One 8:e70962.

Seong E, Yuan L, Arikkath J (2015) Cadherins and catenins in dendrite and synapse morphogenesis. Cell Adh Migr 9:202–213.

Shimoyama Y, Tsujimoto G, Kitajima M, Natori M (2000) Identification of three human type-II classic cadherins and frequent heterophilic interactions between different subclasses of type-II classic cadherins. Biochem J 349:159–167.

Song JY, Ichtchenko K, Südhof TC, Brose N (1999) Neuroligin 1 is a postsynaptic cell-adhesion molecule of excitatory synapses. Proc Natl Acad Sci U S A 96:1100–1105.

Sparrow SS, Cicchetti VD, Balla AD (2005) Vineland adaptive behavior scales, 2nd edition. Circle Pines, MN: American Guidance Service

Stan A, Pielarski KN, Brigadski T, Wittenmayer N, Fedorchenko O, Gohla A, Lessmann V, Dresbach T, Gottmann K (2010) Essential cooperation of N-cadherin and neuroligin-1 in the transsynaptic control of vesicle accumulation. Proc Natl Acad Sci U S A 107:11116–11121.

Suzuki SC, Inoue T, Kimura Y, Tanaka T, Takeichi M (1997) Neuronal circuits are subdivided by differential expression of type-II classic cadherins in postnatal mouse brains. Mol Cell Neurosci 9:433–447.

Takeichi M, Abe K (2005) Synaptic contact dynamics controlled by cadherin and catenins. Trends Cell Biol 15:216–221.

Tripodi M, Evers JF, Mauss A, Bate M, Landgraf M (2008) Structural homeostasis: compensatory adjustments of dendritic arbor geometry in response to variations of synaptic input. PLoS Biol, 6:e260.

van Harssel JJ et al. (2013) Clinical and genetic aspects of PCDH19-related epilepsy syndromes and the possible role of PCDH19 mutations in males with autism spectrum disorders. Neurogenetics 14:23–34.

Voronin LL, Cherubini E (2004) ’Deaf, mute and whispering’ silent synapses: their role in synaptic plasticity. J Physiol, 557:3–12.

Wang C, Pan YH, Wang Y, Blatt G, Yuan XB (2019) Segregated expressions of autism risk genes Cdh11 and Cdh9 in autism-relevant regions of developing cerebellum. Mol Brain 12:40.

Wang K et al. (2009) Common genetic variants on 5p14.1 associate with autism spectrum disorders. Nature 459:528–533.

Willemsen MH, Fernandez BA, Bacino CA, Gerkes E, de Brouwer AP, Pfundt R, Sikkema-Raddatz B, Scherer SW, Marshall CR, Potocki L, van Bokhoven H, Kleefstra T (2010) Identification of ANKRD11 and ZNF778 as candidate genes for autism and variable cognitive impairment in the novel 16q24.3 microdeletion syndrome. Eur J Hum Genet 18:429–435.

Wu N, Wang Y, Pan YH, Yuan X (2020) Identification of CDH11 as an ASD risk gene by matched-gene co-expression analysis and mouse behavioral studies. bioRxiv doi: 10.1101/2020.02.04.931121

Yamagata M, Duan X, Sanes JR (2018) Cadherins Interact With Synaptic Organizers to Promote Synaptic Differentiation. Front Mol Neurosci 11:142.

