## Supplementary figures and images for "Regulation of Neural Circuit Development by Cadherin-11 Provides Implications for Autism"

### Extended Figure 1-1

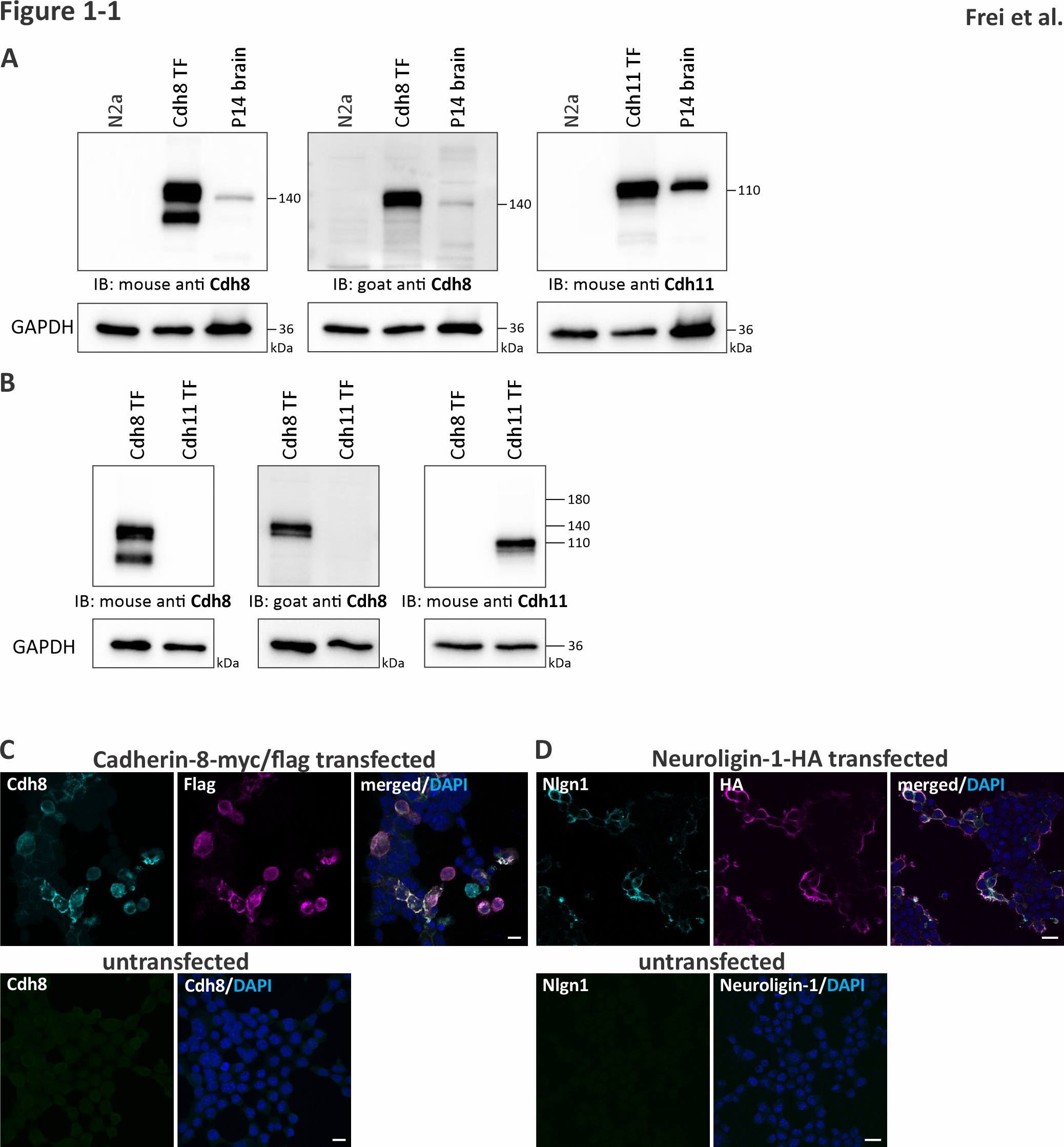
